# *ndufs2^-/-^* zebrafish have impaired survival, neuromuscular activity, morphology, and one-carbon metabolism treatable with folic acid

**DOI:** 10.1101/2025.07.16.664929

**Authors:** Dana V. Mitchell, Donna M. Iadarola, Neal D. Mathew, Kelsey Keith, Christoph Seiler, Sanghyeon Yu, Man S. Kim, Niki Woodard, Vernon E. Anderson, Eiko Nakamaru-Ogiso, Deanne M. Taylor, Marni J. Falk

## Abstract

Mitochondrial complex I (CI) deficiency represents a common biochemical pathophysiology underlying Leigh syndrome spectrum (LSS), manifesting with progressive multi-system dysfunction, lactic acidemia, and early mortality. To facilitate mechanistic studies and rigorous screening of therapeutic candidates for CI deficient LSS, we used CRISPR/Cas9 to generate an *ndufs2^-/-^* 16 bp deletion zebrafish strain*. ndufs2^-/-^* larvae exhibit markedly reduced survival, severe neuromuscular dysfunction including impaired swimming capacity, multiple morphologic malformations, reduced growth, hepatomegaly, uninflated swim bladder, yolk retention, small intestines, and small eyes and pupils with abnormal retinal ganglion cell layer. Transcriptome profiling of *ndufs2^-/-^* larvae revealed dysregulation of the electron transport chain, TCA cycle, fatty acid beta-oxidation, and one-carbon metabolism. Similar transcriptomic profiles were observed in *ndufs2^-/-^* missense mutant *C. elegans* (*gas-1(fc21)*) and two human CI-disease fibroblast cell lines stressed in galactose media. *ndufs2^-/-^* zebrafish had 80% reduced CI enzyme activity. Unbiased metabolomic profiling showed increased lactate, TCA cycle intermediates, and acyl-carnitine species. One-carbon metabolism associated pathway alterations appear to contribute to CI disease pathophysiology, as folic acid treatment rescued the growth defect and hepatomegaly in *ndufs2^-/-^*larvae. Overall, *ndufs2^-/-^* zebrafish recapitulate severe CI deficiency, complex metabolic pathophysiology, and relevant LSS neuromuscular and survival phenotypes, enabling future translational studies of therapeutic candidates.

## INTRODUCTION

Primary mitochondrial diseases (PMD) are a heterogeneous group of disorders that affect at least 1 in 4,300 individuals across all ages and ethnic backgrounds (1). PMD are caused by pathogenic variants in more than 400 genes located in the mitochondrial or nuclear genomes, with extremely variable individual molecular etiologies and phenotypic presentations (2). Clinical manifestations may range from exercise intolerance, weakness, and fatigue to developmental delays, complex neurologic dysfunction, gastrointestinal difficulties, multi-system failure, and death (3). Despite the wide range of symptomatic presentations and prevalence as the most common inborn error of metabolism in the general population, there remain few therapeutic options for patients. Although recent scientific advancements have successfully identified potential therapies for the indication of PMD, demonstrating the efficacy of these candidates remains difficult (4–10).

A common manifestation of PMD in children is Leigh Syndrome Spectrum (LSS) (11), a highly heterogeneous condition with more than 113 associated causal genes (12), which frequently results from pathogenic variants in subunits or assembly factors for complex I (CI, or NADH:ubiquinone oxidoreductase, OMIM 252010) in the mitochondrial electron transport chain (ETC) (12–16). CI plays major roles in oxidative phosphorylation (OXPHOS), both by oxidizing NADH to NAD^+^ and by pumping protons from the matrix to the intermembrane space (IMS) to maintain the proton motive force that is converted to biochemically useful energy as ATP. Indeed, oxidizing much of the NADH generated from the TCA cycle makes CI essential in cellular metabolism for regulating the NADH:NAD^+^ redox balance and maintaining the NAD^+^ pool (17). CI dysfunction thus often disrupts broader cellular metabolism by limiting ATP production and dysregulating oxidative metabolism (18). Additionally, CI is a major contributor to ROS generation, where CI dysfunction may accelerate superoxide production and overwhelm oxidative stress responses (19–22). Disruption of this multitude of essential CI functions contributes to progressive cellular and organ level pathophysiology in LSS individuals who harbor pathogenic variants that alter CI structure, assembly, and function.

Preclinical models offer the opportunity to advance mechanistic understanding of LSS disease pathophysiology and candidate treatments (23, 24). Mouse models of CI deficient LSS often are embryonic lethal or do not display disease phenotypes due to engagement of compensatory pathways not present in humans and are therefore not conducive to screening (25, 26). Two notable exceptions are an *ndufs4,* a knockout model that has provided important mechanistic and therapeutic insights (27) and an ndufs2-null model that recapitulates metabolic alterations typical of LSS (25, 28, 29). However, these models are not amenable to high-throughput treatment screens. Increasingly, the utility of zebrafish (*Danio rerio*) vertebrate animal models for LSS is being demonstrated to facilitate preclinical investigations including high-throughput treatment screens due to their highly conserved nuclear and mitochondrial genetics, rapid development, ease of genetic modification, low cost of maintenance, large brood sizes, and larval stage body transparency that allow *in vivo* examination of system-wide or tissue-specific molecular dysfunction (24, 30). Our group and others have recently established and described several zebrafish genetic PMD models including *ndufaf2^-/-^* (Complex I assembly factor), *surf1^-/-^* (Complex IV assembly factor), *fbxl4^-/-^*(ubiquitin ligase that suppresses mitophagy), *lrpprc^-/-^* (mRNA translation and polyadenylation factor), and *dldh^-/-^* (dihydrolipoamide dehydrogenase), which recapitulate key biochemical and phenotypic aspects of human pathologies (4, 31–34). While a clear advantage of zebrafish models over lower organisms for multi-system disease modeling is their discrete organs, genome and transcriptome sequencing have not yet been widely performed in zebrafish PMD models to inform individual organism or organ level expression changes that may highlight novel aspects of disease pathophysiology and treatment targets.

Here, we report extensive phenotypic and multi-omic characterization of, to our knowledge, the first CI structural subunit mutant zebrafish strain established to better understand LSS organ-level pathophysiology and enable future high-throughput identification of therapeutic candidates for CI disease. Specifically, we used CRISPR/Cas9 to generate zebrafish larvae with a 16 bp deletion from exon 8 in *ndufs2* (NADH dehydrogenase [ubiquinone] iron-sulfur protein 2), encoding one of the core structural components of CI that is a definitive cause of LSS (35–40). In addition to its known role in CI PMD, *ndufs2* in zebrafish is homologous to the *C. elegans* gene *gas-1* (K09A9.5). The *gas-1(fc21)* worms, which are CI-deficient due to a homozygous missense mutation, have been studied extensively over the past two decades. This invertebrate model has provided key insights into their cellular pathophysiology and potential therapeutic targets of PMD (18, 20–22, 41, 42). Because disruption of *gas-1* reliably recapitulates CI deficiency in *C. elegans*, we reasoned that targeting its zebrafish homolog, *ndufs2*, would produce a comparable phenotype, while enabling investigation of tissue- and system-specific effects in a more physiologically complex vertebrate model. Transcriptomic sequencing was performed on our zebrafish model to evaluate the gene expression profile and enable metabolic flux capacity modeling of CI deficiency effects, with results compared to published transcriptome datasets for *C. elegans* CI deficiency *gas-1(fc21)* model and to unpublished internal human patient NDUFS1 and NDUFS4 fibroblast cell lines from our research clinic (17, 20). In addition, unbiased metabolomic studies were performed to evaluate the broader biochemical characteristics of the *ndufs2^-/-^* zebrafish model (See Experimental Schema Overview, **Figure 1**). Results suggest that our CI deficient zebrafish model recapitulates both physical and multi-omic phenotypes of LSS and can therefore be a useful tool for rapid and effective drug screening for LSS and broader PMD therapy development.

**Figure 1:**
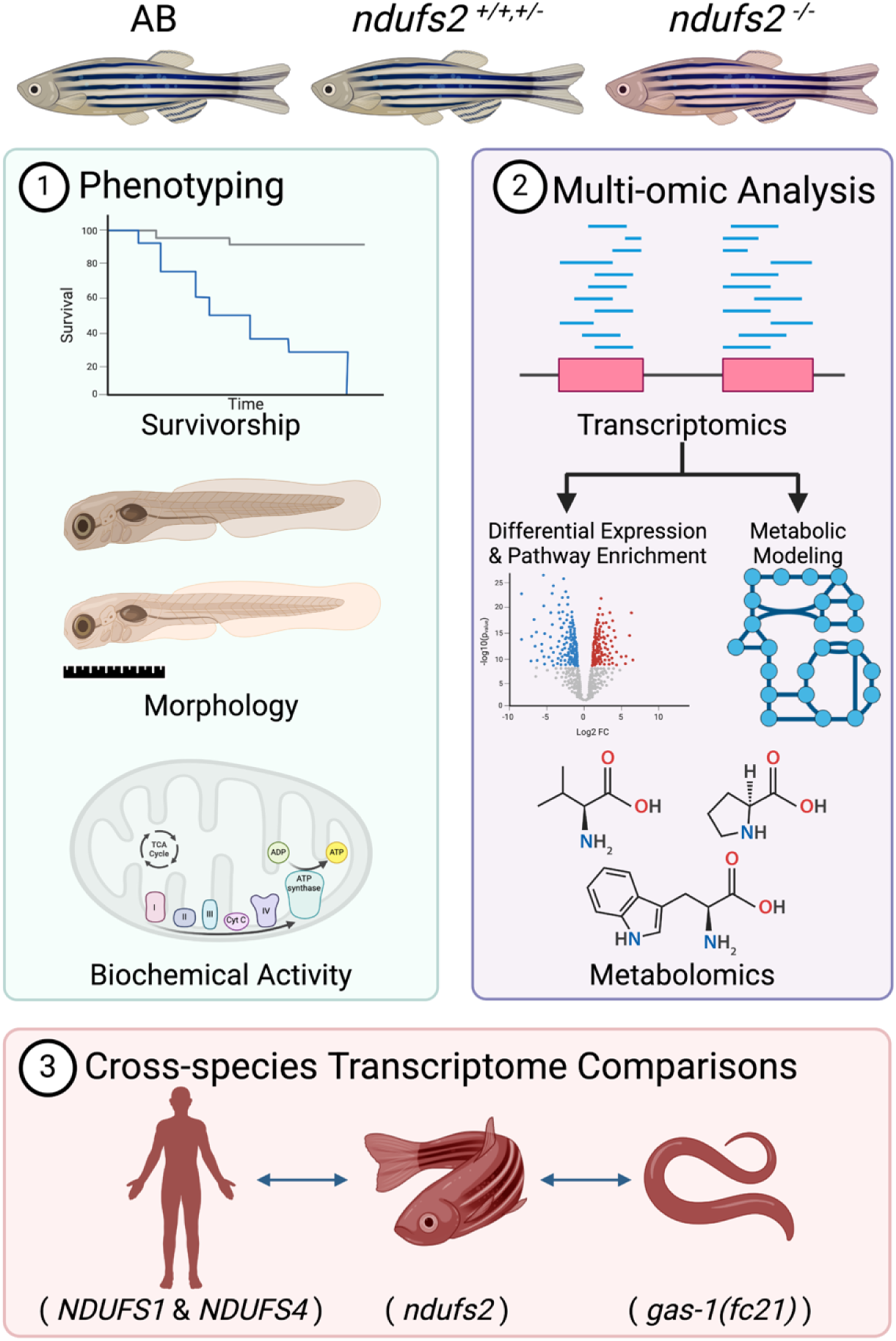
Experimental Schema Overview. Experiments were performed to (1) Characterize a novel CI zebrafish model of LSS generated with CRISPR/Cas9 that harbors a homozygous 16 bp deletion in *ndufs2* using survival, morphologic, and targeted biochemical measurements; (2) Investigate the transcriptome and metabolome of *ndufs2^-/-^* zebrafish relative to wild-type controls; and (3) Compare RNA-Seq transcriptome profiles from the mutant zebrafish with *C. elegans* (*gas-1*(*fc21*) worms with a homozygous missense mutation in NDUFS2) and primary fibroblast cell lines from two human patients with genetic-based CI deficiency (NDUFS1 and NDUFS4) to investigate the broad cellular pathophysiology and validate the comparative utility of this zebrafish model for future studies of CI disorders.

## RESULTS

### Establishment and phenotypic characterization of a novel *ndufs2^-/-^*knockout zebrafish strain

NDUFS2, a catalytic core subunit of CI of the ETC, is highly conserved between *C. elegans*, *D. rerio*, and humans. The amino acid identity between zebrafish and humans is 87% (**Supplemental Figure S1**). To model human CI deficiency, we generated a stable genetic zebrafish model of NDUFS2 deficiency using CRISPR/Cas9 technology (**Figure 2A**). The *ndufs2* deletion zebrafish line was generated by injection of a sgRNA targeting the *ndufs2* sequence, gttcttccagatacgattatTGG, into AB wild-type (WT) zebrafish. The resultant *ndufs2^-/-^* deletion line was obtained by heterozygote incrossing and confirmed by Sanger sequencing, which identified the presence of a 16 bp deletion in exon 8 causing a frameshift starting at amino acid 286 and a premature stop codon at amino acid 313 in homozygous larvae that results in truncation of *ndufs2*. ETC complex enzymatic activities quantified by spectrophotometry confirmed the *ndufs2 ^-/-^* larvae had a significant deficiency of CI enzyme activity, with a mean decrease of 80% relative to WT controls while there was no significant decrease observed in complex II, complex IV or citrate synthase (**Figure 2B**). In addition, we measured and compared NAD^+^ and NADH levels in *ndufs2^-/-^* larvae compared with healthy siblings (*ndufs2^+/+,+/-^*) to determine the downstream effects of dysregulated complex I activity (**Supplemental Methods**). NAD^+^ levels in *ndufs2^-/-^* larvae were significantly decreased to 62.6% of healthy siblings, while NADH levels in ndufs2^-/-^ zebrafish were elevated by 2.17-fold compared to healthy larvae (**Supplemental Figure S2**). Thus, this *ndufs2 ^-/-^* zebrafish mutant has a specific and substantial CI deficiency with overall metabolic consequences.

**Figure 2:**
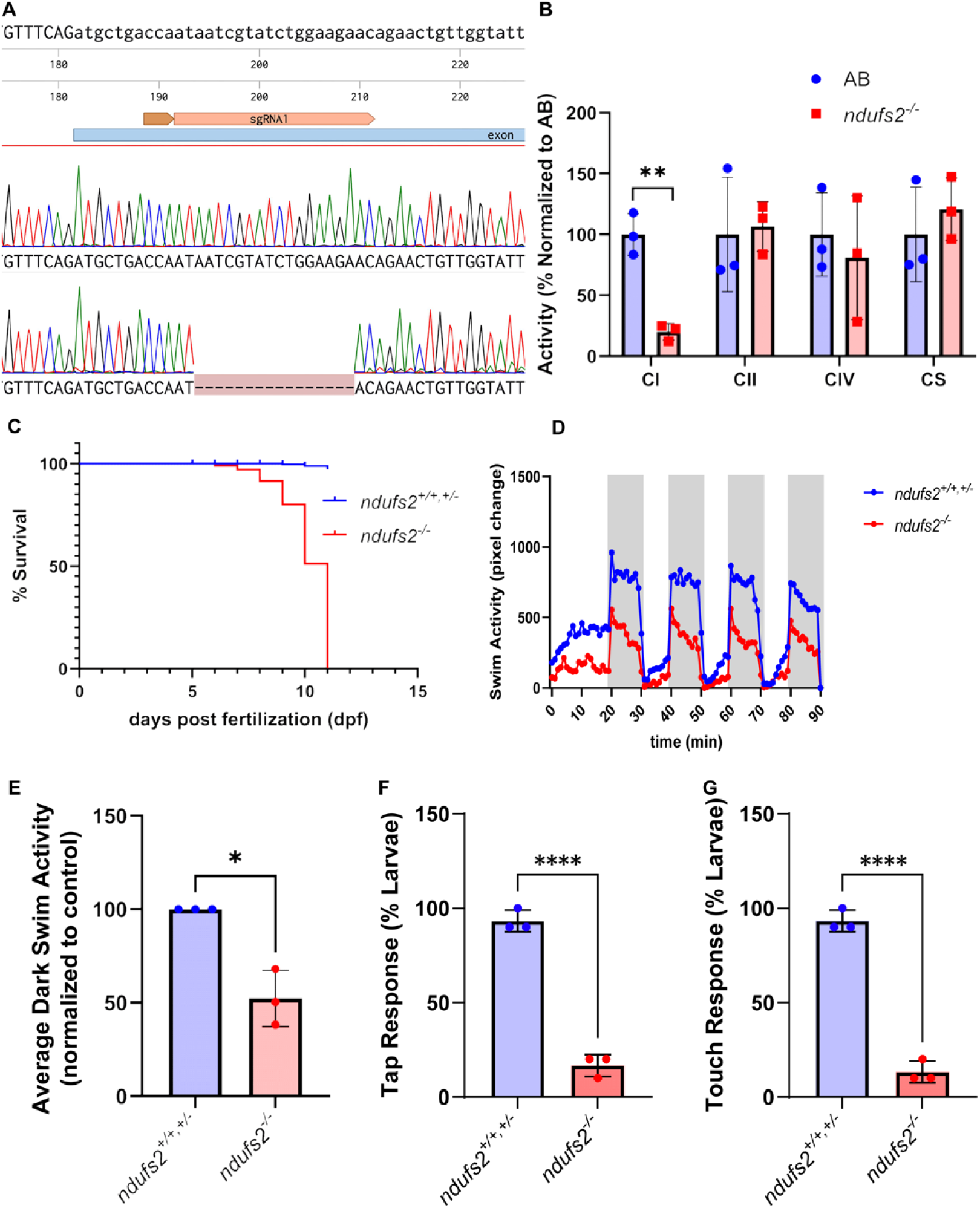
*ndufs2^-/-^*larval zebrafish have isolated CI enzyme deficiency, decreased survival, and decreased swim activity. A) DNA sequence showing the targeted window of *ndufs2* by sgRNA for Crispr editing. Electropherogram confirmation in WT zebrafish of sequencing of this region and the corresponding 16 base pair deletion created in exon 8 of *ndufs2^-/-^*larvae. B) CI, CII, CIV, and citrate synthase (CS) enzyme activities were quantified by spectrophotometric analysis in 7 dpf larvae (n=3). Statistical significance between groups of larvae was determined using a two-tailed Welch’s t-test, where **, p < 0.01. C) Kaplan-Meier curve showing reduced survival of *ndufs2^-/-^* larvae relative to mixed *ndufs2^+/+^*and *ndufs2^+/-^* wild-type sibling larvae (100 larvae from a *ndufs2^+/-^* incross, n=3 biological replicate experiments were performed totaling 300 larvae). The median survival is at 10 dpf and the max survival is at 11 dpf. D) Swim activity in light-dark cycles was quantified in 7 dpf larvae using the Zebrabox (12 larvae per experimental trial, with n=3 biological replicates performed). Dark periods are shaded as grey. E) Quantification of the average swim activity across four dark cycles for all 12 larvae per condition in each experimental trial. Each point represents the average of each biological replicate (n=3). Neuromuscular defect was quantified using F) touch and G) tap startle response of biological replicates (n=3). Statistical significance between groups of larvae for all experiments was determined using two-tailed, Welch’s t-tests (*, p < 0.05; **, p < 0.01; ***, p < 0.001; ****, p < 0.0001).

Next, we characterized the survival and swimming-based neuromuscular activity of the *ndufs2^-/-^* larvae. The *ndufs2^-/-^*larvae showed significantly decreased survival relative to WT siblings with a 50% probability of survival at 10 dpf and 100% mortality at 11 dpf (**Figure 2C**). By comparison, WT zebrafish (AB or *ndufs2^+/-^*) live into adulthood, typically beyond 2 years. Neuromuscular function at the level of swimming activity was quantified using a Zebrabox system. Both *ndufs2^-/-^*larvae and WT siblings responded to light/dark transitions and exhibited increased swim activity during dark cycles (**Figure 2D**). However, *ndufs2^-/-^* larvae showed a significant (p < 0.05) decrease in swim activity during dark cycles of about 50% mean swim activity, indicative of reduced neuromuscular function (**Figure 2E**). To further quantify neuromuscular dysfunction, we performed an assessment of evoked escape response using both touch and tap stimuli. *ndufs2^-/-^*larvae displayed a significant decrease in both touch and tap responses compared to healthy *ndufs2^+/+,-/-^* larvae, providing additional support for neuromuscular defects in the mutant larvae (**Figure 2F & 2G**).

Comparing the gross morphology of *ndufs2^-/-^* larvae with WT siblings at 5 dpf revealed an uninflated swim bladder that remained throughout their lifespan. *ndufs2^-/-^* larvae at 5 dpf also appeared to have an enlarged and darker liver (suggestive of hepatomegaly) and smaller, darker intestines. The enlarged yolk mass in *ndufs2^-/-^*larvae is indicative of slower yolk absorption (**Figure 3A & 3B**). Notable differences in size of the eye and pupil were observed, both appearing smaller in *ndufs2^-/-^* larvae (**Figure 3C & 3D**). Image observations were confirmed with morphological measurements showing *ndufs2^-/-^*larvae had larger and darker livers (**Figure 3E & 3F**), larger yolk mass (**Figure 3G**), smaller eyes and pupils (**Figure 3H & 3I**), and smaller and darker intestines (**Figure 3J & 3K**). *ndufs2^-/-^* larvae size was also significantly smaller by 5% (p < 0.01) in overall animal length measured from nose to tail at 5 dpf (**Figure 3L**). Reassessment of morphological parameters at 7 dpf was performed, revealing similar findings of smaller eyes, smaller intestine, and increased yolk mass in *ndufs2^-/-^* larvae; however, relative differences between *ndufs2^-/-^*and WT larvae in body length as well as in liver size and intensity were diminished or no longer present (**Supplemental Figure S3**). At 7 dpf, there is no longer a significant difference in liver darkness between WT and *ndufs2^-/-^*larvae, which is not unexpected, as unfed 7 dpf WT larvae commonly develop darker livers. Interestingly, histological hematoxylin and eosin staining in 7 dpf larvae show an altered retinal ganglion cell layer of the eye in *ndufs2^-/-^* larvae, where this layer is more granular in WT larvae compared to the denser appearance of the *ndufs2^-/-^* larvae (**Supplemental Figure S4**). Overall, *ndufs2^-/-^* larvae demonstrated multiple physiological abnormalities including a lifespan blunted before maturity, reduced swimming activity indicative of impaired neuromuscular function, and gross morphological abnormalities including linear growth defect, hepatomegaly, uninflated swim bladder, yolk retention, small intestinal size, and small eye and pupil size with an abnormal retinal ganglion cell layer.

**Figure 3:**
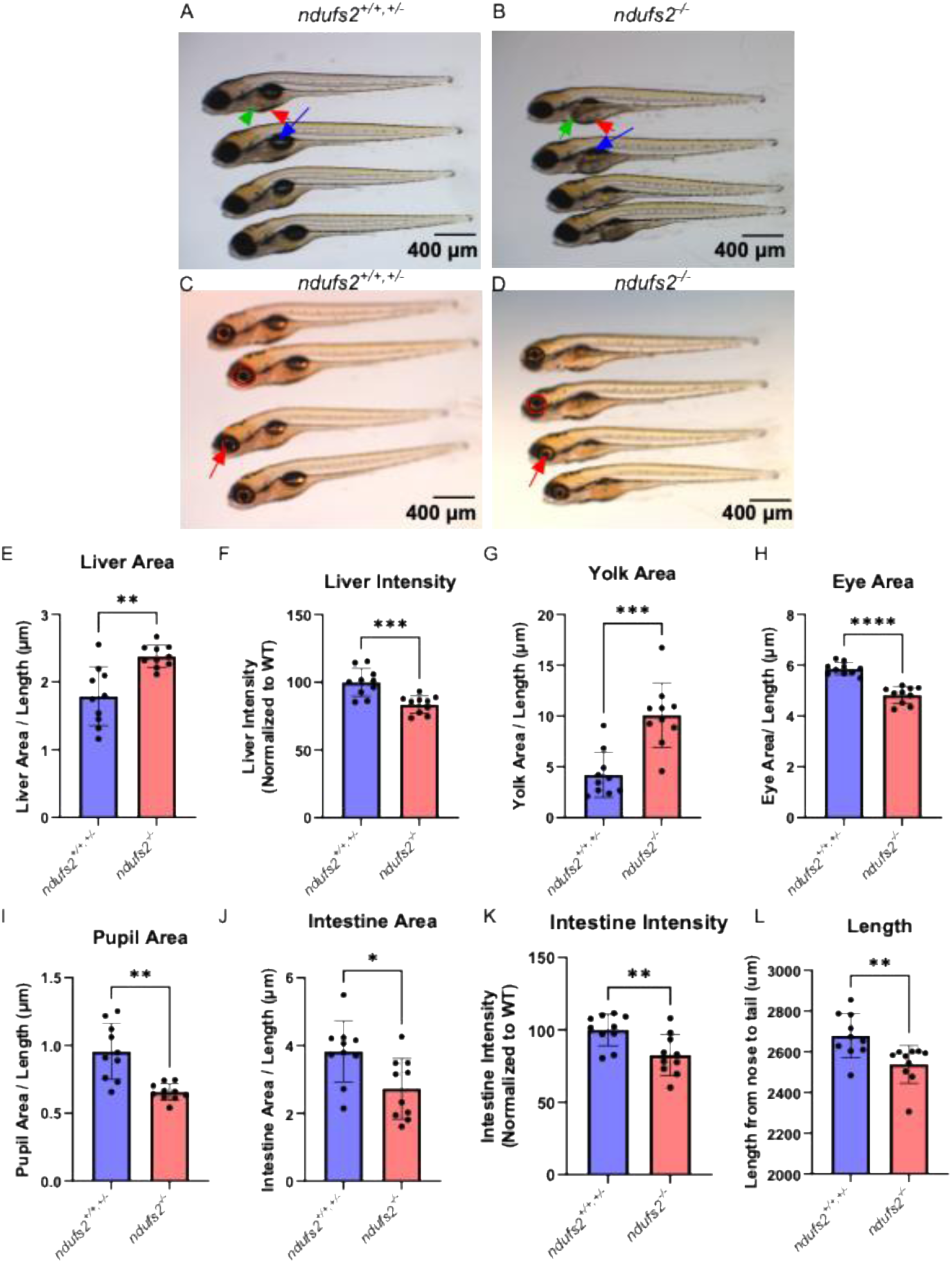
*ndufs2^-/-^ zebrafish* larvae display differences in swim bladder, liver, yolk, eye, and pupil morphology. Light microscope images of 5 dpf larvae demonstrate differences highlighted by arrows between (A) *ndufs2^+/+,+/-^* wild-type and (B) *ndufs2^-/-^* larvae in appearance of their swim bladder (blue), liver (green), and yolk (red). Light microscope images from overhead light show differences between (C) *ndufs2^+/+,+/-^* and (D) *ndufs2^-/-^* larvae in their eye (red circle) and pupil (red arrow) morphology. Black scale bars in each panel represent 400 µm. Quantification was performed on 5 dpf organ-level anatomy and intensity, as well as whole animal length, where differences between larval groups were identified using a two-tailed Welch’s t-test (*, p < 0.05; **, p < 0.01; ***, p < 0.001; ****, p < 0.0001). Measurements include (E) liver area; (F) liver mean intensity; (G) yolk area; (H) eye area; (I) pupil area; (J) intestine area; (K) intestine mean intensity; and (L) length from nose to tail (n=10 total larvae from 3 biological experiments). Smaller intensity reflects an opaquer organ which indicates the image is darker. Similar morphologic differences were detected at 7 dpf, except for normalized hepatic mass (see **Supplemental Figure S2**).

### Transcriptomic Profiling Analyses

#### Differential expression analysis (DEA)

RNA-Seq was performed on *ndufs2^-/-^* mutant zebrafish larvae to establish a transcriptomic profile of the CI deficiency disorder and evaluate cellular pathophysiology at the level of gene and pathway expression differences relative to healthy controls. First, the potential impact on gene expression of experimental variables (or batches) such as clutch, age, and genotype of the zebrafish larvae controls were evaluated (**Supplemental Table S1**). In the absence of previous publications detailing transcriptomic profiling in zebrafish models of PMD and outlining potential sources of variation and error, we compared: (1) clutch effect by comparing expression in offspring from different mating pairs; (2) age effect by comparing 6 dpf and 7 dpf larvae; and (3) genotype of controls effect by testing both AB and mixed population *ndusf2^+/+,+/-^* siblings that were phenotypically normal. Batch effect impact on gene expression was evaluated by performing principal components analysis (PCA) with transcriptomic data from all samples. The transcriptomic profile of samples by PCA were no more similar within than across batches, indicating that clutch, age (6 vs 7 dpf), and genotype of healthy zebrafish larvae (AB and *ndufs2^+/+,+/-^*) have little impact on gene expression (**Supplemental Figure S5**). We also assessed batch effects by quantifying the number of differentially expressed genes in numerous batch comparisons (**Supplemental Figure S6A**). Batch effects due to clutch, age, and control genotypes had little impact on the transcriptomic profile, as shown by low proportions of differentially expressed genes (DEGs) when comparing batch conditions (i.e., 6 vs 7 dpf) in contrast to the number of DEGs identified when comparing *ndufs2^-/-^* larvae to healthy larvae, *ndufs2^+/+,+/-^* or WT (**Supplemental Figure S6A – S6H**). Collectively, these data demonstrate that (1) larvae do not need to be from the same clutch, (2) AB and *ndufs2^+/+,+/-^*siblings can be used interchangeably as healthy controls, and (3) larvae can be collected at either 6 or 7 dpf for transcriptomic studies.

Distinct transcriptomic profiles were identified between *ndufs2^-/-^*larvae and healthy controls (*ndufs2^+/+,+/-^*) (**Figure 4A**). DEA revealed 3,375 genes (13.2%) with significant differential expression (adjusted p-value < 0.05) and an absolute value log2 fold change > 0.58 (**Supplemental Figure S6B**). As expected, *ndufs2* expression showed a significant 96% mean decrease in mutants relative to WT controls (**Figure 4B**). A preponderance of mitochondrial metabolic genes encoding enzymes required for the ETC (*atp5mc1*, *cox6b1*), fatty acid beta-oxidation (*acaa2*, *acsbg1*, *cpt1ab*, *cpt1ab*, *cpt1b*, *cpt2*, *eci2*, *hadhaa*, *hadhab*), monounsaturated fatty acid synthesis (*scd*), oxidoreductase activity (*sesn2*), and mitochondrial OXPHOS uncoupling (*ucp2*) were significantly upregulated in *ndufs2^-/-^* larvae compared to healthy controls (**Figure 4C**). Genes involved in nucleotide metabolism and other one-carbon metabolism associated pathways were also significantly upregulated in *ndufs2^-/-^*larvae, including genes associated with the distribution of one-carbon groups (*aldh1l2*), purine biosynthesis (*atic, mthfd1l*, *paics*, *ppat*), pyrimidine synthesis (*cad*, *upp1*), DNA replication and maintenance (*cdt1*, *gins2*, *mcm3*, *mcm4*, *orc3*, *orc4*, *orc6*), nucleotide formation and metabolism (*dut*, *tyms*), glutathione synthesis (*gss*), methylation (*mat1a*), and methionine synthase (*mtr*) (**Figure 4C**). Genes associated with eye development and visual activities were significantly downregulated, including those related to crystallin (*cryaa*, *crybb2*), retina development (*dnase1l1l*, *fgf19*), lens development and maintenance (*hsf4*, *lgsn*, *mipa*, *mipb*), and visual perception (*kera*) (**Figure 4C**).

**Figure 4:**
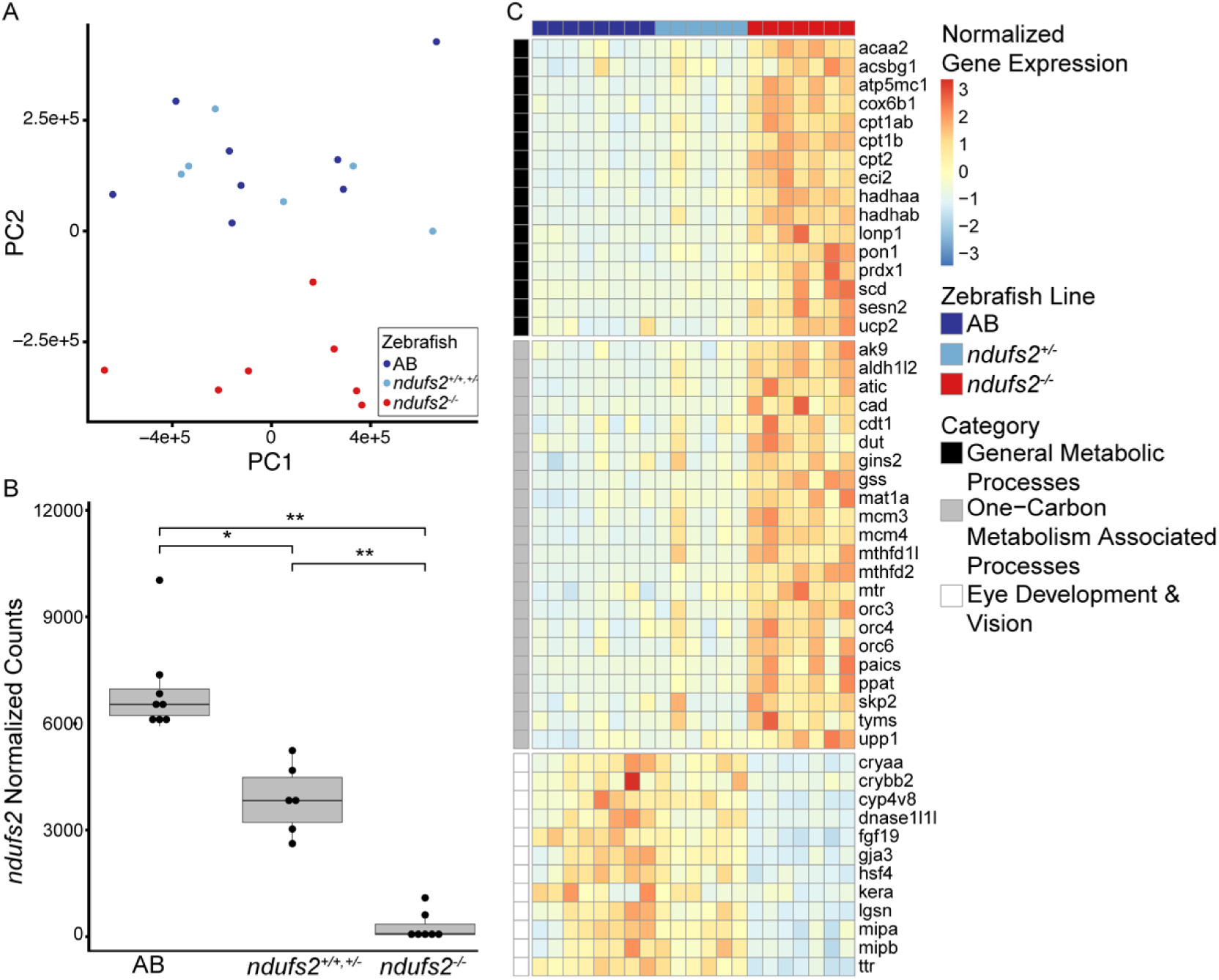
Transcriptomic gene-level analysis of *ndufs2^-/-^* larvae. A) Principal component analysis plot from RNA-Seq data indicating the zebrafish line of the larvae. Each dot represents a single fish colored according to genotype. AB fish are wild-type with genotype *ndufs2^+/+^* and marked in dark blue. In cross siblings with mixed genotypes *ndufs2^+/+,+/-^* are marked in light blue. Mutant *ndufs2^-/-^* zebrafish are marked in red. See **Supplemental Figure S4** for PCA coded by days post fertilization and clutch. B) Normalized transcript counts to quantify *ndufs2* gene expression. Differences in normalized counts between groups were assessed using an unpaired, two-tailed Student’s t-test with Bonferroni correction for multiple comparisons (* p < 0.001, ** p < 0.0001). Significant differences were seen in *ndufs2* normalized counts between all groups: AB and *ndufs2^+/+,+/-^* (44% decrease in *ndufs2^+/+,+/-^*, P = 0.0003); AB and *ndufs2^-/-^* (96% decrease in *ndufs2^-/-^*, p < 0.0001); and *ndufs2^+/+,+/-^* and *ndufs2^-/-^* (92% decrease in *ndufs2^-/-^*, p = 0.0001). C) Heatmap of differentially expressed genes in *ndufs2^-/-^* compared to AB and *ndufs2^+/+,+/-^*larvae. Gene expression is presented as normalized transcript counts and scaled across genes. All genes represented have significant differences in expression between groups (p < 0.05) calculated using DESeq2 (v 1.38.3). Genes are grouped based on zebrafish pathway categories labeled on figure for horizontal groupings.

#### Gene set enrichment analysis (GSEA)

To understand the pathways most impacted by differences in gene expression, we performed GSEA using GeneOntology (43), Reactome (44), Kyoto Encyclopedia of Genes and Genomes (KEGG) (45), and WikiPathways (46) zebrafish ontological databases through the R package WebGestaltR (47).

Several pathways were significantly upregulated, as indicated by their normalized enrichment score (NES) in *ndufs2^-/-^* larvae compared with healthy controls. Upregulated pathways included those associated with general mitochondrial and metabolic processes such as the ETC, fatty acid metabolism and TCA cycle (**Figure 5**), consistent and expected with mitochondrial dysfunction (17, 48–50). One-carbon metabolism and nucleotide metabolism pathways were also enriched in *ndufs2^-/-^* larvae (**Figure 5**). Alternatively, healthy zebrafish larvae were enriched compared with *ndufs2^-/-^*larvae for pathways involved in eye/lens development such as lens development in camera-type eye (**Figure 5**). These findings are consistent with the morphological analysis reported above that confirm *ndufs2^-/-^* larvae have an eye-specific developmental defect. Overall, transcriptomic profiling validates use of the *ndufs2^-/-^*larvae model through identification of expected gene and pathway expression changes that provide insights to other understudied pathways that contribute to the complex pathophysiology of CI disease.

**Figure 5:**
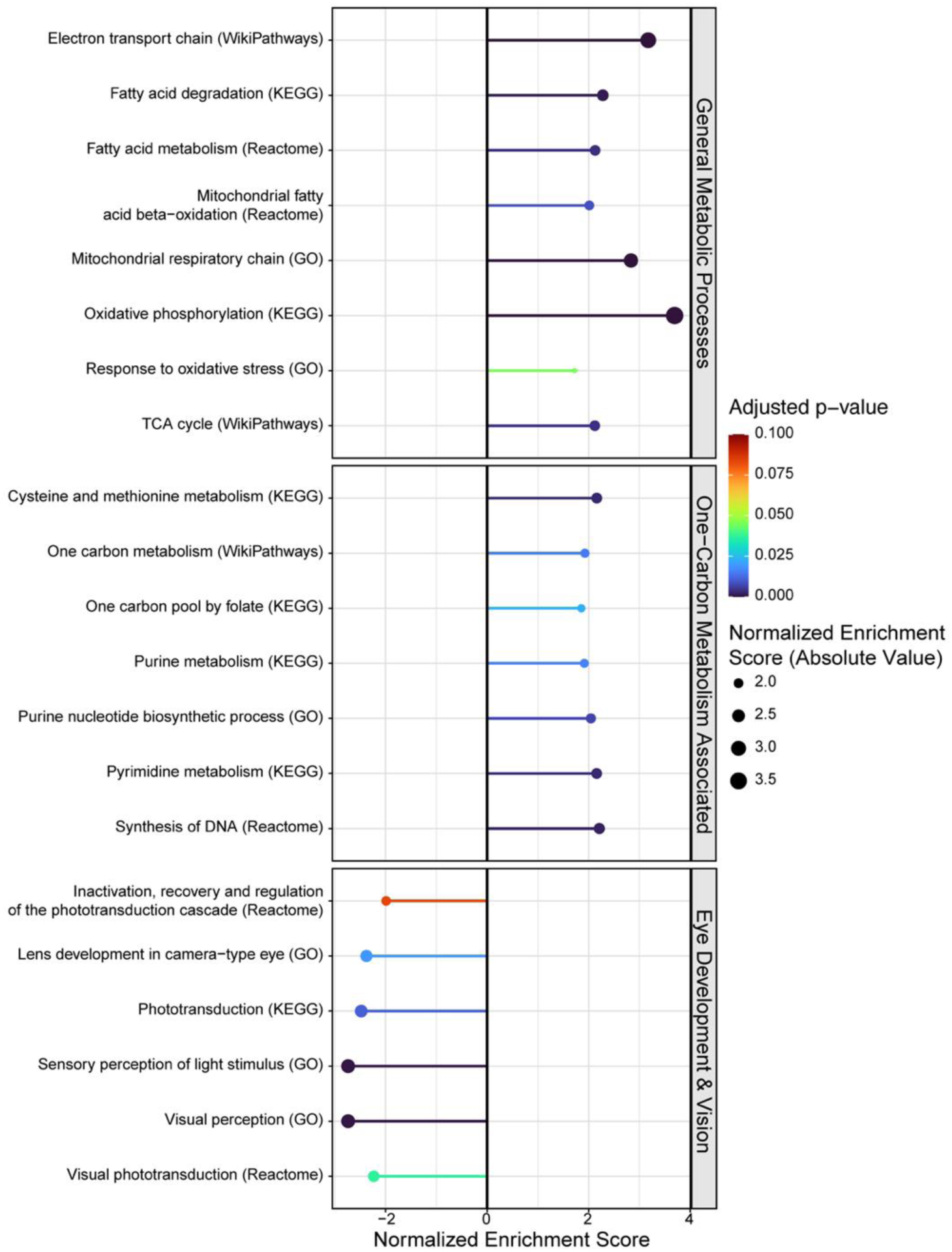
Transcriptomic pathway analysis of *ndufs2^-/-^* larvae. *ndufs2^-/-^* larvae show enrichment of expression in pathways related to general metabolic processes and one-carbon and nucleotide metabolism but downregulated expression of pathways involved in eye development and vision. Pathway enrichment analysis was performed with RNA-Seq data using GO, KEGG, and WikiPathways databases in *ndufs2^-/-^* larvae compared to healthy control larvae through the R package WebGestaltR. Pathways with lollipops towards the right or left indicate pathways that are upregulated or downregulated, respectively, in *ndufs2^-/-^* larvae relative to wild-type controls.

#### Metabolic flux modeling

Using flux-balance analysis (FBA) applied to our zebrafish transcriptomic data, we simulated flux through a comprehensive zebrafish genome-scale metabolic model (GEM), Zebrafish1 (51), to determine if there were predicted differences in the activity of biochemical reactions in *ndufs2^-/-^* compared to *ndufs2^+/+,+/-^*larvae. FBA revealed increased flux capacity in *ndufs2^-/-^* compared to healthy larvae of reactions comprising general metabolic processes. Specifically, predicted dysregulated metabolic reactions included those upregulated in the pentose phosphate pathway such as the phosphoribosyl pyrophosphate synthetase (PRPPS) Zebrafish1 reaction (7% increase in *ndufs2^-/-^* larvae), and two transketolase Zebrafish1 reactions (TKT1 & TKT2) (5% increase in each) (**Supplemental Figure S7, Supplemental Table S2**). Reactions in the mitochondrial beta-oxidation of even-chain fatty acids were predicted to have an overall 54% increase of flux capacity in *ndufs2^-/-^* larvae. More generally, the fatty acid oxidation (FAO) system showed increased predicted metabolic flux capacity such as a 100% increase for the reaction converting 3-hydroxyisovaleryl-CoA to 3-hydroxyisovaleryl-carnitine (FAOXC5OHc, p < 0.01) and a 100% increase in the reaction transporting 3-hydroxyisovaleryl-CoA from the mitochondria into the cytosol (HIVCACBP, p < 0.01) (**Supplemental Figure S8A & S8B**). Genes previously identified to have significantly increased expression in *ndufs2^-/-^* larvae participate in these metabolic reactions such as *acsbg1*, *cpt1ab*, and *eci2* in FAO and more specifically, genes in the subsystem beta-oxidation of even-chain fatty acids such as *hadhaa*, *hadhab*, and *acaa2* (**Figure 4C**).

Biochemical processes involved in one-carbon metabolism associated activities were also predicted to have increased biochemical flux capacity in *ndufs2^-/-^*larvae such as folate metabolism with an overall 25% increase in predicted flux capacity and purine metabolism with large increases in predicted flux capacity of specific reactions such as the adenosine kinase reaction (ADNK1, 86% increase, p < 0.01) (**Supplemental Figure S8C & S8D**).

### Cross-species transcriptome profiling comparison with the *ndufs2^-/-^* zebrafish model

#### Human CI disease patient fibroblast cell lines

We performed a comparative analysis of the *ndufs2^-/-^*zebrafish transcriptomic profile with that of unpublished fibroblast cell lines from human patients with similar gene disorders involving CI subunits (NDUFS1 and NDUFS4) to further evaluate the use of *ndufs2* mutant zebrafish as a proxy for understanding the pathophysiology of CI mitochondrial disorders in humans. Human cell lines were grown in galactose media to induce metabolic stress conditions and reveal any transcriptome adaptations that are masked when cultured in glucose (52). After mapping zebrafish genes to their human orthologs, gene to gene comparisons showed that zebrafish normalized average expression counts per million (cpm) were highly correlated with averaged CI human cell line normalized average expression cpm (r = 0.39, p < 0.001) (**Figure 6A**). Concordant gene expression changes confirm that the *ndufs2^-/-^*zebrafish CI disease model has utility to inform cellular pathophysiology, and potentially the efficacy of therapeutic candidates for human PMD when studied in zebrafish models. We also identified an overlap in genes that were significantly upregulated in *ndufs2^-/-^* larvae compared with healthy controls and humans with CI disorders compared to healthy human-derived fibroblast cell lines (n = 7, **Figure 6B**). These included genes encoding mitochondrial proteins essential for proper mitochondrial function (G0S2, *HSPD1,* and *MPV17L*) and genes implicated in eye disorders as were also prominent in the previously described GSEA of *ndufs2^-/-^* zebrafish larvae (*LAMA1*) (**Figure 6C**).

**Figure 6:**
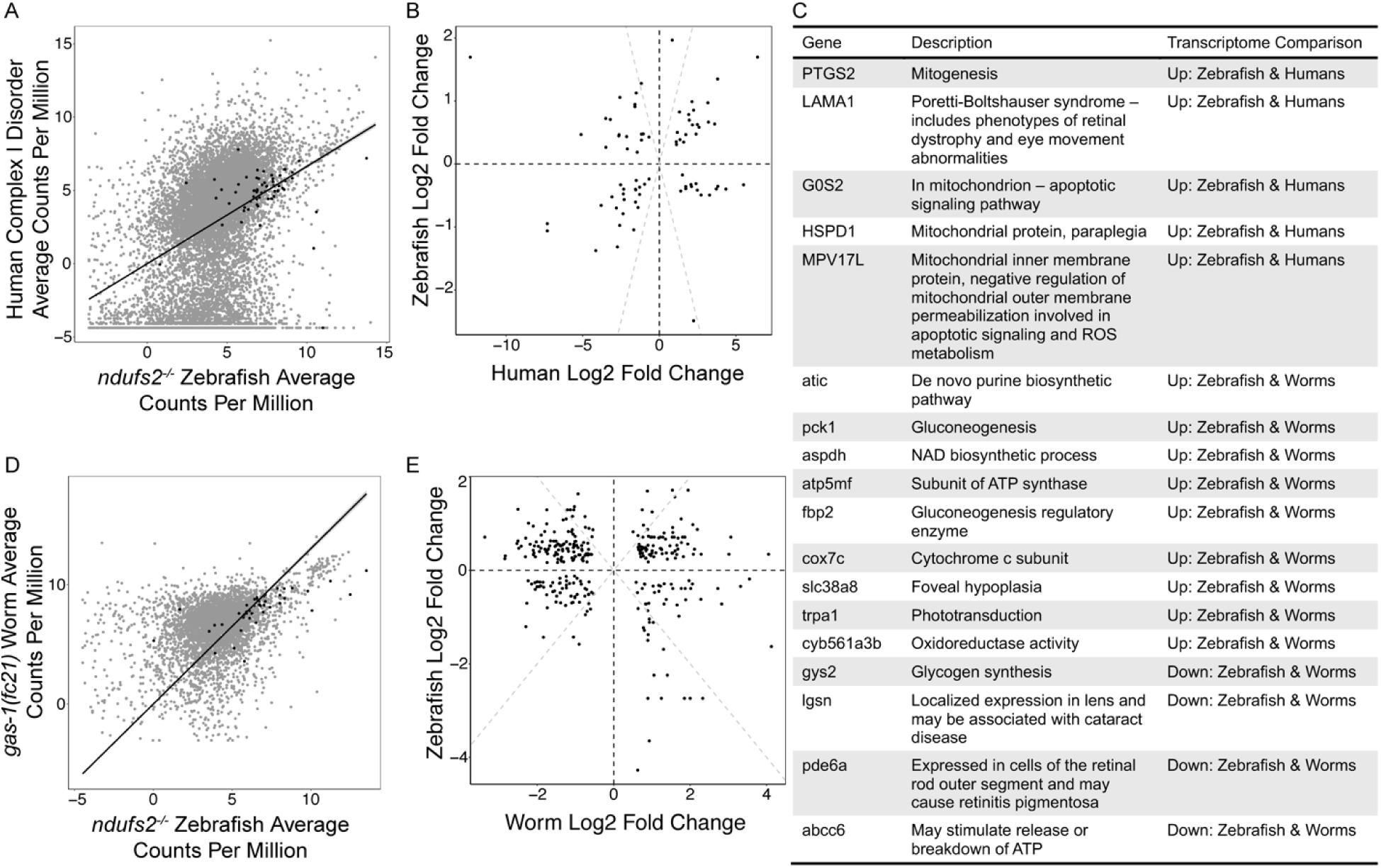
Cross-species comparisons of transcriptomic data in nuclear gene encoded CI structural subunit disorders. A) Comparison of normalized transcript counts in *ndufs2^-/-^* larvae with human CI disease fibroblast cell lines (n=2 individuals, *NDUFS1^-/-^* and *NDUFS4^-/-^*). Overall gene expression was highly correlated between humans and zebrafish (Pearson correlation, r = 0.39, p < 0.001). ETC genes are highlighted in black and line of best fit extending through the origin is displayed. B) Comparison of normalized transcript counts in *ndufs2^-/-^* larvae with *gas-1(fc21)* worms (n=4). Overall gene expression was highly correlated between worms and zebrafish (Pearson correlation, r = 0.70, p < 0.001). ETC genes are highlighted in black and line of best fit extending through the origin is displayed. C) Comparison of log2 fold change of differentially expressed genes (adjusted p-value < 0.01 calculated using DESeq2) between human CI disease fibroblast cell lines and *ndufs2^-/-^* zebrafish. The 1:1 relationship in each direction is shown with the black dotted line. D) Comparison of log2 fold change of differentially expressed genes (adjusted p-value < 0.01 calculated using DESeq2) between *gas-1(fc21)* worms and *ndufs2^-/-^* zebrafish. The 1:1 relationship in each direction is shown with the black dotted line. E) Table of metabolic related genes with similar patterns of differential expression across model organism transcriptome profiles. The description column gives a short explanation of the relevance of each gene. The Transcriptome Comparison column states whether the gene is upregulated or downregulated and in which organisms.

#### gas-1(fc21) Caenorhabditis elegans

Similarities in gene expression profiles were identified between *ndufs2^-/-^*larvae and *gas-1(fc21)* worms. *gas-1(fc21)* is a well-established *C. elegans* strain containing a homozygous p.R290K missense mutation in the CI subunit homolog of NDUFS2 (53), which has served as an effective invertebrate model in which to characterize the organismal effects and pathophysiology of CI deficiency (18, 20–22, 41, 42). After mapping worm genes to their zebrafish orthologs, overall normalized average expression cpm were highly correlated between the two animal models (r = 0.39, p < 0.001). Genes encoding proteins of the ETC had an even stronger correlation between *ndufs2^-/-^* larvae and *gas-1(fc21)* worms (r = 0.70, p < 0.001) (**Figure 6D**). This relationship suggests that the *ndufs2^-/-^* CI zebrafish model is comparable to other standard models of mitochondrial CI disease.

Genes were identified that had similar patterns of differential expression in *ndufs2^-/-^* zebrafish larvae compared with *ndufs2^+/+,+/-^* larvae as did *gas-1(fc21)* worms compared with N2 wild-type worms (**Figure 6E**). Genes that had significantly increased expression in both zebrafish and worms with CI disorders included those directly encoding mitochondrial proteins or ETC subunits (*hspa4*, *hsdl2*, *atp5mf*, and *cox7c*), involved in purine biosynthesis (*atic*), and related to vision or sight (*slc38a8* and *trpa1*), among others. An overlap of genes having significantly decreased expression was found in both CI disease model organism comparisons for vision-related disorders or cells of the eye (*lgsn* and *pde6a*, respectively) (**Figure 6C**).

### Unbiased metabolomics profiling

We sought to develop unbiased metabolomics in zebrafish larvae to independently validate transcriptomic findings. Indeed, partial least squares discriminant analysis (PLS-DA) (54) recapitulated the results of our transcriptomic analysis by showing tight biological replicates and a clear separation of wild-type AB larvae from *ndufs2^-/-^* larvae (**Supplemental Figure S9A**). Variable importance projections (VIP) of PLS-DA showed that tissue lactate, the most common blood biomarker elevated in PMD, is among those metabolites with the highest differences between *ndufs2^-/-^* and healthy larvae. Additionally, two TCA cycle intermediates, fumarate and malate, have high VIP scores, validating the expected TCA cycle disruption from ETC disfunction. Multiple *N*-acetyl amino acids and acyl-carnitines were also included among the metabolites with the highest VIP scores, indicating an increase in the presence of acyl-CoAs from FAO (**Figure 7A**). These metabolites are noted to be present in various genetic diseases that also disrupt FAO (55) and are used for the diagnosis of human PMD of FAO (56, 57). Metabolomics pathway enrichment analysis indicates that *ndufs2^-/-^* larvae are enriched compared with healthy larvae for one-carbon metabolism pathways. Other enriched metabolic pathways in *ndufs2^-/-^* larvae included pantothenate and CoA biosynthesis, as expected from the increase in acyl-carnitines (58), and pyrimidine metabolism (**Supplemental Figure S9B**).

**Figure 7:**
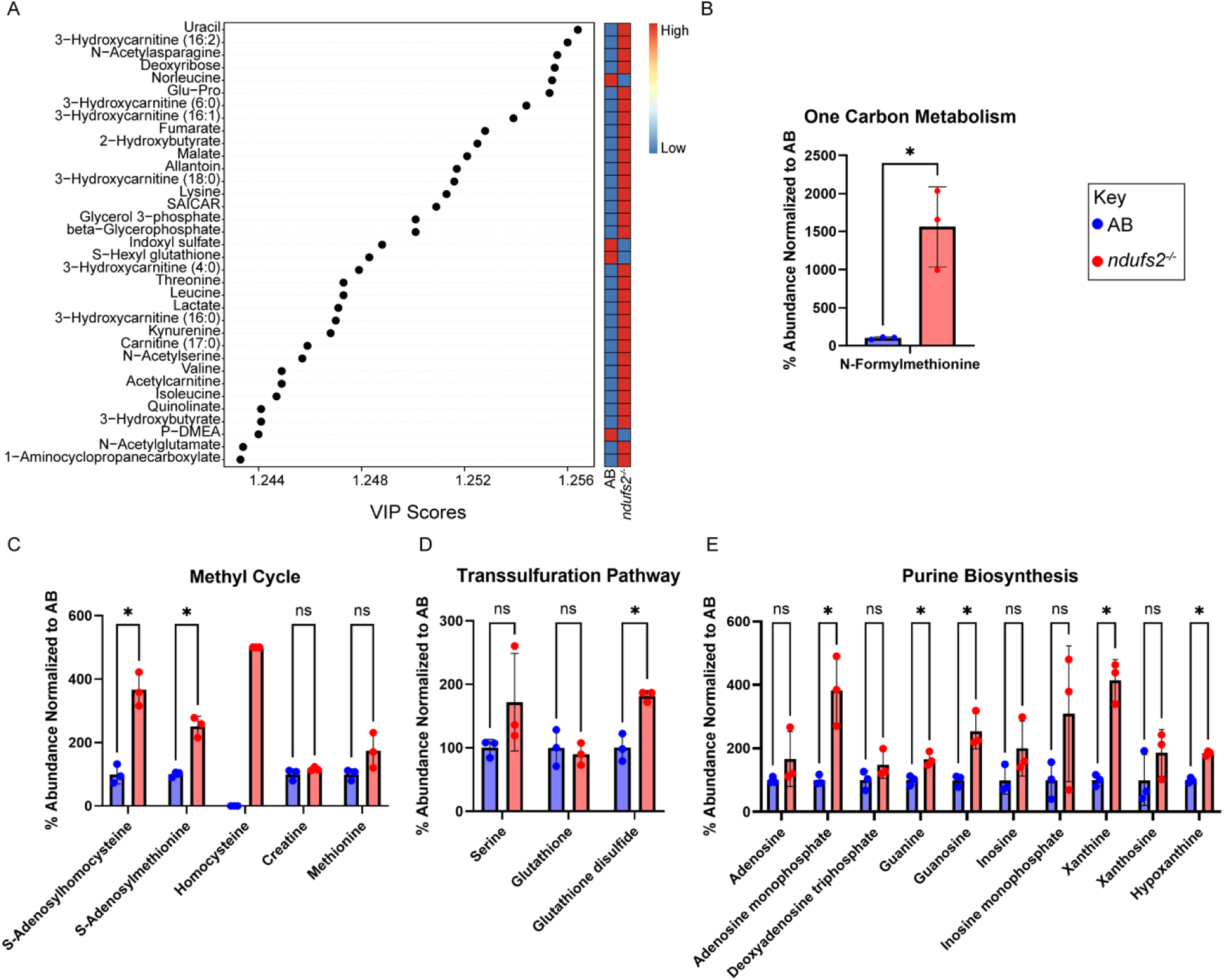
Untargeted metabolomics analysis shows upregulated one-carbon metabolism and one-carbon associated pathways in *ndufs2^-/-^* zebrafish larvae. A) Variable importance projection (VIP) of the top 30 metabolites significantly up or downregulated in *ndufs2^-/-^* larvae compared only to AB (p < 0.05). Metabolites are shown on the y-axis and the x-axis shows the VIP score, a measure of the contribution the variable makes towards the PLS-DA (**Supplemental Figure S7A**). The heatmap to the right indicates directionality of the change in metabolite by showing the average metabolite levels of AB and *ndufs2^-/-^* zebrafish, respectively. B) *N-*-formylmethionine metabolite levels as a marker for the one-carbon metabolism pathway (n=3 biological replicates each containing 20 larvae). C) The methyl cycle is also linked to one-carbon metabolism, with 5 identified metabolites quantified (n=3 biological replicates each containing 20 larvae). As homocysteine was not detected in AB fish, it could not be normalized to AB but instead is arbitrarily shown as *ndufs2^-/-^* larvae having 500% enrichment compared to AB. D) Three metabolites were identified in the transsulfuration pathway, which is an associated pathway of one-carbon metabolism, of which glutathione disulfide was significantly increased in *ndufs2^-/-^*larvae (p < 0.05, n=3 biological replicates each containing 20 larvae). E) Purine biosynthetic pathway is linked to one-carbon metabolism, of which many intermediates and metabolites were identified in this dataset (n=3 biological replicates each containing 20 larvae). Statistics were calculated using a two-tailed Welch’s t-test: ns, not significant; * p < 0.05.

RNA-Seq transcriptome profiling identified many significantly dysregulated metabolic pathways involving the same metabolites whose levels were significantly dysregulated in our unbiased metabolomics profiling dataset of *ndufs2^-/-^* larvae. For example, glutathione metabolism was upregulated in the *ndufs2^-/-^*transcriptome and metabolomics profiling showed *ndufs2^-/-^* larvae had a significant increase in levels of glutathione disulfide. This provides an example of transcriptional upregulation that effectively results in an elevation of the expected metabolite. In contrast, branch chain amino acid (BCAA) degradation was upregulated in the transcriptome consistent with changes observed in prior CI disease models (50), while metabolomics showed clear elevations in the BCAA levels of valine, isoleucine, and leucine, indicating less BCAA catabolism (**Figure 7A**). This result demonstrates a transcriptional adaptation that is responding to a likely primary metabolic problem that cannot be fixed at the transcriptional level.

Mitochondrial DNA (mtDNA) replication disorders have recently been recognized to cause dNTP imbalance and changes in the one-carbon cycle (59). Although CI deficiency has not been previously linked to mtDNA replication defects, CI dysfunction is a likely sequela downstream of impaired mtDNA metabolism since the mtDNA encodes 7 core CI structural subunits. Further, as detailed above, significant disruption of one-carbon metabolism and nucleotide metabolism is evident in the *ndufs2^-/-^*transcriptomic dataset. We therefore searched our metabolomics dataset to confirm that one-carbon metabolism is disrupted at the metabolite level. *N-*formylmethionine (fMet), a metabolite in one-carbon metabolism, was identified as one of the most elevated metabolites with 145% change (p = 0.04) (**Figure 7B & 9**), but no other metabolites present in the metabolomics profiling directly linked to the one-carbon pathway were found to be dysregulated. To further investigate one-carbon metabolism disruption, we looked at related pathways that utilize formate intermediates such as the methylation cycle, transsulfuration, and purine biosynthesis. The methylation cycle showed significant elevations in *S-* adenosylhomocysteine, *S*-adenosylmethionine (SAM), and homocysteine (**Figure 7C & 9**). A review of transcriptomic profiling further supports increased activity of SAM in the methyl cycle by revealing upregulation of the *coq3* (p < 0.001, log2FC = 0.31) and *coq5* genes (p < 0.001, log2FC = 0.54), two genes known to be associated with reactions involving SAM in the mitochondria according to the Zebrafish1 GEM (51) (**Figure 9**). In the transsulfuration pathway, no significant increase was seen in levels of glutathione, but significant increase was seen in levels of glutathione disulfide (**Figure 7D**).

Within the purine biosynthetic pathway, elevations were seen of guanosine, xanthosine, xanthine, hypoxanthine, guanine, and AMP, all indicative of nucleotide imbalance (**Figure 9**). In *ndufs2^-/-^* larvae, expression of *acp6*, a gene involved in reactions of the purine synthesis system that include AMP (51), is significantly upregulated (p = 0.01, log2FC = 0.67), providing evidence for nucleotide imbalance beginning at the transcriptomic level (**Figure 9**). Interestingly, adenosine and inosine levels were not significantly changed, indicating that all purines were not affected equally (**Figure 7E**). These data support *ndufs2^-/-^*larvae having significant disruption in one-carbon metabolism and one-carbon-linked pathways, including transsulfuration, the methylation cycle, and nucleotide metabolism (**Figure 9**). This combined study of transcriptomics and metabolomics in *ndufs2^-/-^* zebrafish larvae demonstrated that the CI deficiency results in significant cell-wide metabolic disruption.

### Folic acid treatment rescues growth defect and hepatomegaly in *ndufs2^-/-^* larval zebrafish

To probe if the disrupted one-carbon metabolism detected in *ndufs2^-/-^*larvae contributed to their phenotypic abnormalities and therefore could be therapeutically modulated to improve animal health, we supplemented larvae with folic acid. We selected folic acid as a treatment after identifying dysregulation in one-carbon metabolism pathways in our RNA-seq analysis. Given that folate is a key metabolite in one-carbon biochemical reactions and is primarily processed in the liver, we hypothesized that folic acid supplementation could reverse the effects of the gene expression changes and resolve the lipid accumulation observed in the organ. In addition, previous research by Tsun-Hsien Hsiao et al (60) showed that an eye-size defect from one-carbon metabolism disrupted larvae, similar to the eye phenotype of our *ndufs2^-./-^* larvae, can be partially rescued with folic acid supplementation. Therefore, we supplemented larvae with 100 µM folic acid starting at 1 dpf. At 5 dpf, a significant increase in the length of the *ndufs2^-/-^* larvae was seen (**Figure 8A**). No significant change was evident in the size of the eye or pupil when normalized by zebrafish length (**Figure 8B & 8C**). Surprisingly, we observed a full rescue of hepatomegaly, with reduction of liver size at 5 dpf in *ndufs2^-/-^* larvae treated with folic acid (**Figure 8D**).

**Figure 8:**
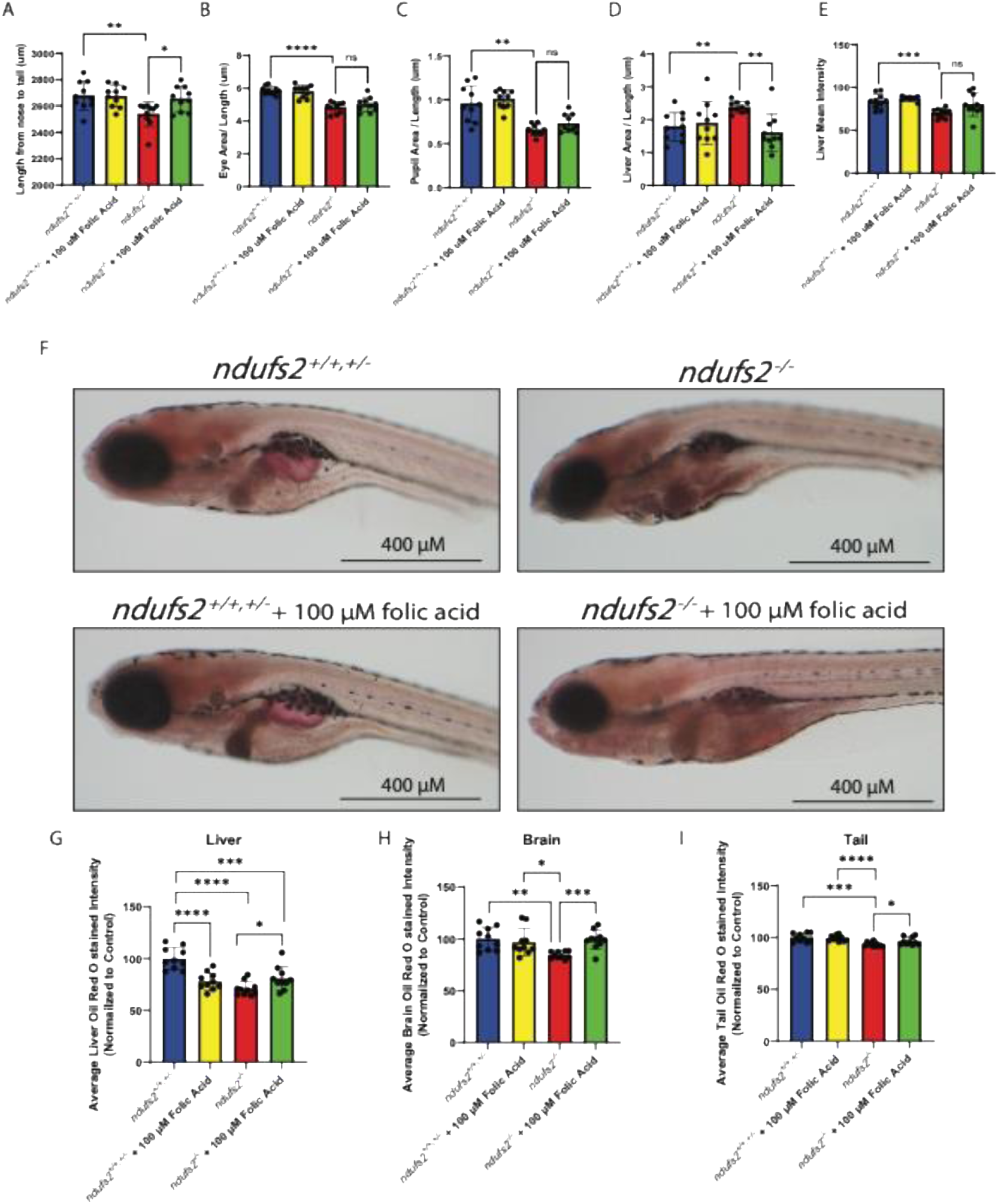
Folic acid supplementation rescued length and liver defects in *ndufs2^-/-^* larvae. Quantification of 5 dpf organs after treatment with folic acid from 1 dpf to 5 dpf: A) length from nose to tail; B) eye area; C) pupil area; D) liver area; E) liver area. F) Representative images from Oil Red O staining of 7 dpf ndufs2+/+,+/- and ndufs2-/-larvae with and without folic acid. Top left, ndufs2+/+,+-/; bottom left, ndufs2+/+,+/ + 100 µM folic acid; top right, ndufs2-/-; bottom right, ndufs2-/- + 100 µM folic acid. Quantification of G) liver, H) brain, and I) tail intensity from Oil Red O staining (n=10). Increased Oil Red O staining is measured by decreased light intensity. Ns, not significant; * p value < 0.05; ** p value < 0.01; *** p value < 0.001; **** p value < 0.0001. P-values were calculated using a two-tailed Welch’s t-test.

The dark liver (**Figure 8E**), enlarged yolk mass, small intestinal size, and impaired swim activity were not rescued by this folic acid treatment (data not shown). Previous PMD zebrafish models have also displayed hepatomegaly and dark livers, which was due to high hepatic lipid and lipid droplets (4, 31, 32). To evaluate if folic acid was contributing to a decrease in liver lipids, we stained whole fixed 7 dpf larvae with Oil Red O (**Figure 8F**). *ndufs2^-/-^* larvae showed increased liver Oil Red O staining (where increased liver staining is measured as decreased light intensity) indicative of increased lipids, consistent with findings in other PMD zebrafish models. Additionally*, ndufs2^-/-^* larvae had a partial rescue in lipid accumulation upon treatment with folic acid (**Figure 8G**). Unexpectedly, more staining in the control larvae were observed with folic acid treatment. Strikingly, folic acid fully rescued high lipids in the brains of *ndufs2^-/-^* larvae (**Figure 8H**). We also measured tail muscle lipid accumulation, where *ndufs2^-/-^* larvae showed a modest increase in lipids that was partially rescued by folic acid treatment (**Figure 8I**). Overall, these findings point towards a specific role of one-carbon metabolism disruption in the impaired animal growth, liver pathology, and lipid metabolism of *ndufs2^-/-^* larvae and the potential of folic acid treatment to improve liver related phenotypes in LSS (**Figure 9**).

**Figure 9:**
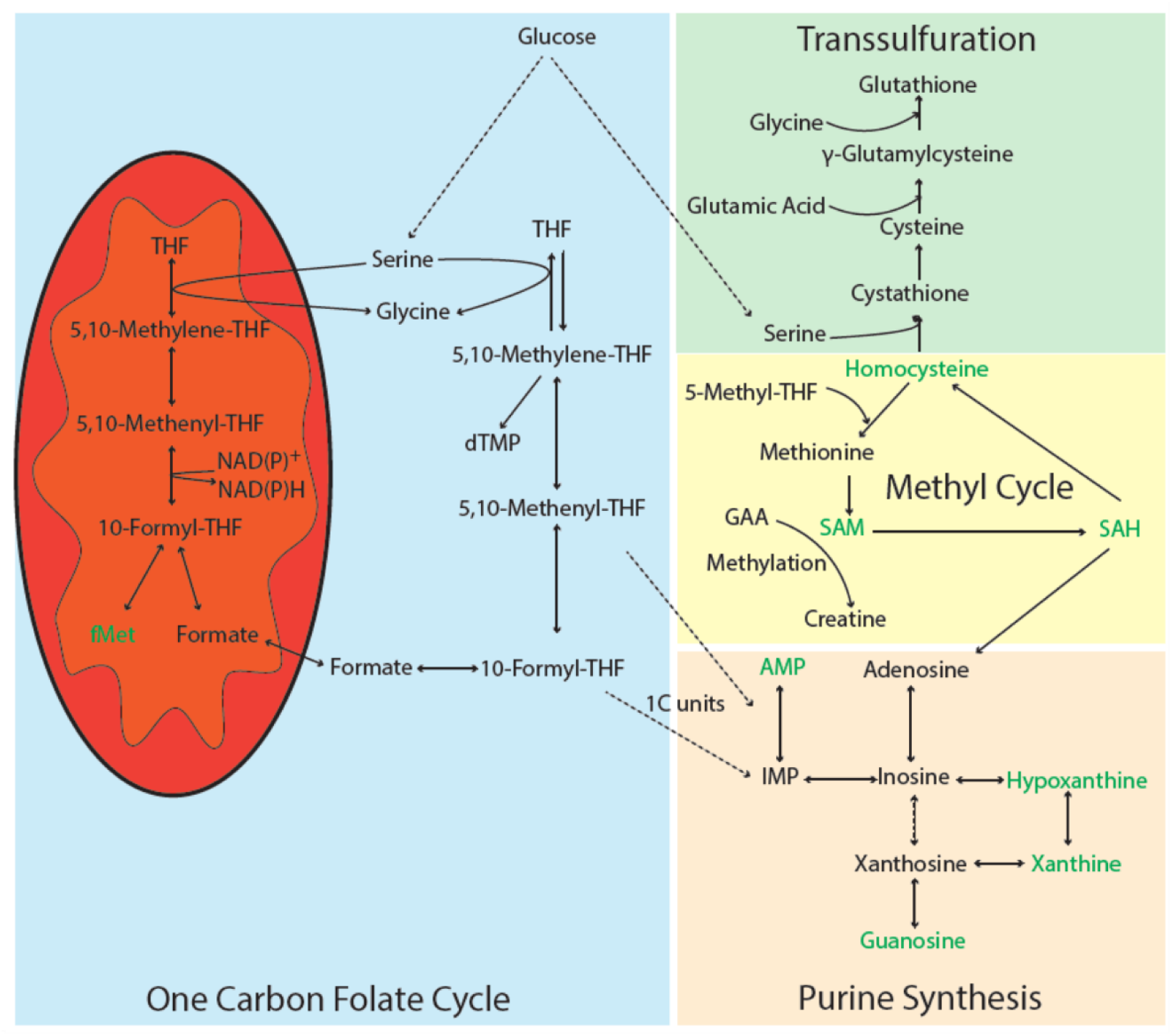
One carbon metabolism and associated pathways (transsulfuration, methyl cycle, and purine synthesis). Green metabolites indicate metabolites that were significantly increased in *ndufs2*^-/-^ larvae. THF, tetrahydrofolate; dTMP, thymidine monophosphate; NAD^+^, nicotinamide adenine dinucleotide; SAH, *S*-adenosyl-homocysteine; SAM, *S*-adenosylmethionine; GAA, guanidinoacetic acid; AMP, adenosine monophosphate; IMP, inosine monophosphate.

## DISCUSSION

In this study, we successfully applied CRISPR/Cas9 technology to generate a stable genetic *ndufs2^-/-^* 16 bp deletion zebrafish line that had 80% reduced CI enzyme activity, 96% reduced *ndufs2* mRNA expression, significantly reduced lifespan (mean 10 dpf, max 11 dpf), swimming-based neuromuscular activity, and multiple gross morphological changes including linear growth defect, hepatomegaly, uninflated swim bladder, yolk retention, small intestinal size, and small eye and pupil size with an abnormal retinal ganglion cell layer. We then developed a unique set of omics-level methodologies including transcriptome and unbiased metabolome sequencing in *D. rerio* (zebrafish) larvae to investigate the complex metabolic dysfunction caused by primary CI deficiency. Specifically, we performed one of the first bulk RNA-Seq studies in a PMD zebrafish model, utilizing the most current zebrafish annotation for analytical quantification to elucidate fine scale changes in gene expression (61) and determining the relative impact of key experimental biological variables. Transcriptome profiling results showed that conditions such as age (6 vs 7 dpf), clutch, and genotype of healthy controls have little effect on gene expression results, providing a baseline for future zebrafish transcriptomic studies, while also unveiling dysfunction in essential metabolic pathways such as the pentose phosphate pathway and FAO. Similarities in gene expression, particularly involving upregulation of ETC, OXPHOS, TCA cycle, and FAO, between the *ndufs2^-/-^* zebrafish model of CI disease when compared with both human fibroblast cell lines from CI disease LSS patients (*NDUFS1^-/-^* and *NDUFS4^-/-^*) and CI deficient worms (*ndufs2^-/-^*, *gas-1*(*fc21*)) demonstrate that the *ndufs2^-/-^* zebrafish model is a useful translational resource for studying CI deficient LSS mitochondrial disorders. Key results from pathway analysis and metabolic flux modeling in transcriptome data were verified with unbiased metabolomics profiling data, yielding new insights into the pathophysiology of CI disorders by their impact on various metabolic pathways such as one-carbon metabolism and nucleotide metabolism (**Figure 9**). Remarkably, treating the zebrafish from 1 to 5 dpf with 100 µM folate rescued their growth defect and hepatomegaly, suggesting a specific role for one-carbon metabolism disruption in the impaired animal growth and liver pathology of *ndufs2^-/-^* larvae.

*ndufs2^-/-^* zebrafish exhibited an enlarged and dark-appearing liver phenotype (**Figure 3A, 3B, 3E & 3F**), similar to previously characterized LSS models (4, 31, 32). Although dark livers are reported to have high lipid and lipid droplet levels, the underlying mechanism(s) is not understood (34).

Hepatomegaly and liver steatosis are pathologies that can be associated with human LSS (62, 63), and appear frequently in larval PMD models (4, 31, 32). We postulate that these liver phenotypes resulted at least in part from disrupted one-carbon metabolism in *ndufs2^-/-^* zebrafish, based on several lines of evidence: (1) We identified several genes and pathways enriched in *ndufs2^-/-^* larvae that related to one-carbon metabolism (**Figure 4C, 5, & 9**); (2) Our genome-scale metabolic flux modeling methodology predicted a system-wide increase in folate metabolism (**Supplemental Figure S7D & Figure 9**); and (3) Unbiased metabolomic profiling confirmed upregulation of these processes with increased abundance in *ndufs2^-/-^*larvae of one-carbon metabolism markers such as fMet (**Figure 7D & 9**). Importantly, a reduction in CI function could conceivably result in disruption of one carbon metabolism due to an impairment in NADH oxidation and an increase in the NADH/NAD+ ratio which regulates methylenetetrahydrofolate dehydrogenase. NAD^+^ and NADH measurements in the zebrafish larvae directly support this hypothesis (**Supplemental Figure S2**). Our metabolomics data identified other pathways also controlled by the NADH/NAD+ ratio such as the TCA cycle and FAO. In addition, previous research has shown that the liver preferentially utilizes NAD(P)H from serine catabolism for lipogenesis, which in turn is increased by the inhibition of CI, CIII, or CV activity (64, 65). Therefore, disruption of CI in *ndufs2^-/-^* larvae may increase serine catabolism, causing excess lipogenesis in the liver. Lipid accumulation in our mutant zebrafish may be exacerbated by impaired fat absorption from the zebrafish yolk in models with mitochondrial dysfunction and is further consistent with dysregulated FAO on transcriptome profiling (**Figure 4 & 5**). Dysregulated lipid metabolism in the liver may also cause the decreased yolk absorption observed in *ndufs2^-/-^* larvae (**Figure 3G**). Folate deficient zebrafish display hepatic lipid accumulation (66), consistent with our hypothesis.

Finally, prevention of enlarged livers with folic acid supplementation in larvae (**Figure 8D, 8F, & 8G**) suggests liver phenotypes result in part from disrupted one-carbon metabolism in the *ndufs2^-/-^*zebrafish model. There is a known albeit weak connection between folate and mitochondrial disease. Specifically, Kearns-Sayre Syndrome (KSS), a PMD caused by a large-scale mtDNA deletion, is associated with low cerebral spinal fluid (CSF), 5-MTHF levels, or folate deficiency (67). Our *ndufs2^-/-^* larvae may display brain phenotypes not typically seen in patients due to their severe CI deficiency and near-complete *ndufs2* depletion by 10–11 dpf. Folinic acid (a natural form of folic acid) treatment has led to neurological improvements in a few patients, similarl to our experimental treatments (67–69). Overall, we demonstrate a connection between liver folate deficiency and lipid accumulation in *ndufs2^-/-^*larvae that is partially rescued by folic acid. Moving forward, we aim to measure folate levels in liver and brain across our PMD models to further validate this link. Future studies that obtain transcriptomic, metabolomic, and methylation data from *ndufs2^-/-^* larvae, both before and after treatment administration, would provide further insight into the underlying mechanisms of our observed morphological and molecular disruptions.

Transcriptome gene, pathway, and metabolic flux capacity profiling similarly highlighted that secondary changes in mitochondrial fatty acid metabolism occur in CI disease. Indeed, CI deficiency introduces an energetic defect while simultaneously increasing the NADH/NAD^+^ ratio, which serves as the driver of FAO. Whereas increased expression of ETC and/or OXPHOS pathways exhibited the greatest change in *ndufs2^-/-^*zebrafish, the second most elevated pathways on transcriptomic profiling involved fatty acid metabolism (**Figure 5**), including carnitine-CoA acyltransferases (*cpt*) and hydroxyacyl-CoA dehydrogenases (*hadha*) (**Figure 4C**). In the absence of external food sources, *ndufs2^-/-^* larvae rely on increased lipid oxidation from their yolk sacks to meet the substantial energy demands for growth. However, the enlarged yolks observed (**Figure 3E**) suggests that these attempts are insufficient. The inability to efficiently utilize lipids from the yolk may be attributed to disruption in lipid metabolism as mentioned previously, an elevated NADH/NAD^+^ ratio inhibiting FAO, and/or GI absorption issues that typically accompany LSS. Transcriptomic analyses reveal upregulation of FAO pathways, indicating a compensatory mechanism as the system attempts to mobilize and utilize yolk lipids.

Furthermore, metabolic flux capacity modeling in the *ndufs2^-/-^*zebrafish also identified increased predicted flux through mitochondrial FAO, highlighting the system’s preparedness to metabolize the required lipids despite the observed inefficiencies (**Supplemental Figure S8B**). Independently, metabolic profiling data further highlighted the disrupted fat metabolism of *ndufs2^-/-^* zebrafish, where 8 of the 30 most enhanced metabolites were acyl-carnitines, including seven 3-hydroxy derivatives (**Figure 7A**) whose levels are expected to be enhanced by increased NADH. In human patients, increased serum and urine acyl-carnitine species are used to identify primary genetic or secondary metabolic defects occurring in mitochondrial FAO, and are used as a proxy for the increased corresponding acyl-CoA species (55, 70–72). Collectively, these data suggest *ndufs2^-/-^* zebrafish have enlarged livers with increased yolk lipid accumulation due in part to impaired FAO flux, owing to their upregulated NADH:NAD^+^ ratio that is caused by their genetically-disrupted CI (NADH dehydrogenase) capacity and was likely exacerbated by disrupted one-carbon metabolism in the liver, as in mouse liver NAD(P)H is derived from serine catabolism (4, 34, 64).

It is interesting to consider whether rescuing the liver lipid phenotype may extend lifespan in *ndufs2^-/-^* CI deficient larvae and other PMD zebrafish models. In the Leucine Rich PPR-motif-Containing protein (LRPPRC) deficient zebrafish model of French Canadian Leigh Syndrome, for example, genetic reintroduction of LRPPRC specifically to the liver significantly improved lifespan (73). However, the therapeutic benefit of folic acid supplementation in the *ndufs2^-/-^*model implies that liver improvement alone may be insufficient to rescue the overall *ndufs2^-/-^* larval multi-system phenotype. Thus, alternative treatments such as a CNS-penetrant form of folinic acid, serine, or methionine may hold potential value for their ability to rescue the broader problems of reduced survival, neuromuscular activity impairment, and multi-organ morphology defects of *ndufs2^-/-^* zebrafish.

A particularly striking morphological observation in our *ndufs2^-/-^* larvae was their significantly smaller eye and pupil size, with an abnormal retinal ganglion cell layer, as compared to WT siblings (**Figure 3C, 3D, 3H & 3I**), which has not been observed in other LSS zebrafish larvae models.

Transcriptomic analysis provided mechanistic insight into this morphological presentation as many genes associated with eye, retina, and/or lens development were downregulated in *ndufs2^-/-^*larvae. Genes associated with eye disorders were also dysregulated such as corneoretinal dystrophy (*cyp4v8*) (74), cataracts (*gja3*, *hsf4*, and *lgsn*) (75–78), and congenital cornea plana (*kera*) (79, 80) (**Figure 4C**). Of particular interest, vision related genes were also downregulated upon our comparative transcriptome analyses in human CI disease fibroblast cell lines and CI disease *C. elegans*, two systems without obvious visual relevance. Decreased activity of lens development, visual perception, and phototransduction pathways in the CI disease zebrafish model suggest that the observed decrease in pupil size results from disruption in lens development at the larval stage, causing visual impairment (**Figure 5**) (81). The small eye defect, or microphthalmia, has also been reported in a zebrafish model of folate deficiency, further corroborating the link between one-carbon metabolism and pathology (60). One-carbon metabolism has been shown to be disrupted in some patients with PMD through cerebral folate deficiency (82). Despite current research suggesting folate derivatives and dNTPs should be investigated as treatments for PMD (60, 83), we were not able to increase eye and pupil size using 100 µM folate (**Figure 8B & 8C**) (64). It remains possible that future studies further modulating the timing, formulation, dosing, and/or ocular penetrance of treatments that replete one-carbon metabolism may improve eye development in *ndufs2^-/-^* larvae.

Transcriptomic and unbiased metabolomic modeling in *ndufs2^-/-^*larvae uncovered a host of dysregulated pathways and genes that may be downstream cellular effects of primary CI deficiency that significantly impact animal survival and health and thereby warrant investigation for potential therapeutic manipulation. Here, we primarily focused on one-carbon metabolism, as multiple lines of evidence showed its clear disruption ranging from gene expression and pathway dysregulation to morphological manifestations in *ndufs2^-/-^*larvae and further supported by recent literature. However, we also observed molecular changes previously reported and commonly observed in other PMD models including CI disruption such as: (1) enrichment in general metabolic pathways like the ETC, OXPHOS, and TCA cycle (**Figure 5**) (84); (2) a system-wide predicted increase in flux capacity through biochemical reactions associated with FAO (**Supplemental Figure S8A & S8B**) (20); and (3) increased metabolite abundance such as glutathione disulfide and lactate (4, 21, 32, 34) (**Figure 7A & 7D**). These adaptive changes typical of PMD further validate our zebrafish model as a useful translational research system in which to study the pathophysiology of specific CI gene mutations. Strong similarities in gene expression profiles between *ndufs2^-/-^*larvae and both human fibroblast and worm CI structural subunit nuclear gene disease models demonstrate the utility of using our model to understand underlying genetic perturbations caused by PMD and the effects of potential therapeutics (**Figure 6**). In addition, the metabolic modeling methods we have detailed here can be adapted to alter pathways hypothesized as targets for LSS treatment, providing a cost-effective, computational experiment for initiating the development of novel therapeutics. A further advantage of the zebrafish model of CI disease is the ability to evaluate a wide variety of organ-level effects, which are not possible to ascertain in cell line or *C. elegans* systems.

## CONCLUSION

Overall, this work represents important advances in the field of zebrafish biomedical research and demonstrates how zebrafish models can be used in a multi-omic framework; creates a vertebrate animal model of severe CI deficiency with reduced survival, neuromuscular activity, and multi-organ involvement; and provides insight into key cellular mechanisms that drive CI phenotypes and may be amenable to targeted treatment. Identification of the therapeutic role of folic acid in reversing growth and liver defects in this CI disease model suggests further investigation of treatments that target defective one-carbon metabolism is warranted in rigorous clinical trials in human mitochondrial CI disease with LSS and potentially broader clinical manifestations of complex I deficient PMD.

## METHODS

### Characterization of a *ndufs2^-/-^* mutant zebrafish line

#### Sex as a biological variable

Sex was not considered as a biological variable in this study due to it being indeterminate in zebrafish juvenile hermaphrodite larvae at 6 and 7 dpf (85, 86).

#### D. rerio strains and maintenance

AB WT zebrafish were obtained from the zebrafish international research center (ZIRC, [https://zebrafish.org/]). The *ndufs2^-/-^* mutant line was generated at the Children’s Hospital of Philadelphia (CHOP) Zebrafish Aquatics Core using CRISPR/Cas9 gene editing. The 16bp deletion allele is designated as cri4 at the Zebrafish Information Network (https://zfin.org). A sgRNA targeting *ndufs2* (gttcttccagatacgattatTGG) was injected into AB WT embryos at the 1-cell stage and animals were grown to adulthood to generate the F0 generation. Offspring of the F0 generation were confirmed by PCR and Sanger sequencing. A 16 bp deletion was confirmed (AATCGTATCTGGAAGA) in exon 8 of *ndufs2* resulting in a frameshift mutation starting at amino acid 286 and a premature stop codon at amino acid 313. As this truncation eliminates the 4Fe-4S binding site (AAs 326-347), it was anticipated to disrupt the activity of CI (Genbank Accession CT583661).

F1 generation adult *ndufs2 ^+/-^*zebrafish were mated to obtain *ndufs2 ^-/-^* mutants and their phenotypically WT siblings of mixed genotypes, delineated *ndufs2 ^+/+,+/-^.* The embryos were collected on 0 dpf and grown in embryo media (E3) at 28℃ with the following composition: 5.0 mM NaCl, 0.17 mM KCl, 0.33 mM CaCl2, 0.33 mM MgSO_4_. The embryos were sanitized with 0.003% sodium hypochlorite on 1 dpf with three alternating washes with water 5 min each. Pronase was used to promote uniform hatching on 1 dpf by addition of 170 µL 20 mg/mL Pronase to 30 mL E3 media for 3-4 hours and transfer pipette mixing to release embryos from chorions (87). E3 media was replaced on 5 dpf with 10 mM Tris E3 media pH 7.2. Homozygous *ndufs2 ^-/-^* mutants were consistently identified by an uninflated swim bladder by 5 dpf. Zebrafish maintenance and husbandry was performed at 28°C in the CHOP Zebrafish Core.

#### Survival analysis

Survival was monitored by placing 100 larvae from an *ndufs2 ^+/-^* incross into a Petri dish at 1 dpf after bleaching and dechorionation. Scoring was performed every day starting at 5 dpf until all homozygous fish were scored as inviable. Fish were fed with GEMMA Micro 75 fish food starting at 7 dpf once daily. Statistical analysis was performed using the log rank (Mantel-Cox) test with Graphpad Prism (Graphpad Software, San Diego, CA). Lifespan analysis was performed with three biological replicates.

#### Larval swimming activity

Swimming activity was assessed by Zebrabox methodology which monitors larval movement in light and dark periods using the Zebrabox tracking system and Zebralab software (ViewPoint Life Sciences; Montreal, Canada), based on previously published protocols (4, 18, 32). Larvae at 7 dpf were placed individually into wells of a square-well 96-well plate with 200 µL of Tris-buffered E3 media and allowed to acclimate to the Zebrabox under 100% light power for 20 min 28°C prior to experimentation. The experiment consisted of four cycles of a 10 min dark period (no light) followed by 10 min of 100% light (1170 lux). Each biological replicate included technical replicates of 12 WT siblings (*ndufs2^+/+,+/-^*) and 12 *ndufs2 ^-/-^* larvae. Three independent biological replicates were performed per condition. The average AUC in each dark period for all replicates was used to analyze the Zebrabox data using Graphpad Prism. We evaluated neuromuscular dysfunction my quantifying startle response of both healthy and *ndufs2^-/-^* zebrafish larvae. We performed this assessment with both touch and tap evoked escape response using biological replicates (n=3).

#### Morphology analysis

Zebrafish larval morphology was assessed using a light dissecting microscope (Olympus MVX10, Center Valley, PA), an Olympus DP73 camera, and Olympus CellSens® imaging software. Pictures of the eyes were obtained by shining an overhead light on the fish in addition to the microscope illumination from below. ImageJ was used to quantify the various parameters (88).

#### ETC enzyme activity assays

ETC enzyme activity assays were performed by spectrophotometry, as previously described (4).

### Transcriptomics

#### Sample preparation

Five pooled zebrafish larvae were flash frozen at 6 or 7 dpf per condition. Zebrafish were collected from 6 varying clutches (**Supplemental Table S1**). Qiagen RNeasy Plus Mini Kit (cat. # 74134) was used for the extraction, following the manufacturer’s protocol. More details on RNA extraction can be found **Supplemental Materials**.

#### Alignment & Quantification

RNA-Seq data were preprocessed using the nf-core RNAseq pipeline version 3.9 (89).

#### Differential expression analysis (DEA)

Following preprocessing, data were filtered using the R package WGCNA (v. 1.69) to remove genes with many missing entries or zero variance (90). Normalization and differential expression between pairs of groups were performed using DESeq2 (v. 1.38.3) (91). Comparisons included: healthy vs. affected homozygous *ndufs2* mutants (overall, at 6 dpf, and at 7 dpf), 6 vs. 7 dpf in each zebrafish line, healthy AB zebrafish vs. *ndufs2^+/+,+/-^* siblings, and similarities between 3 different clutches (**Supplemental Table S1**). These comparisons aimed to understand the impact of the *ndufs2* mutation on gene expression as well as to confirm that other characteristics of the larvae had little effect on transcriptomic profiles. Genes with an adjusted p-value of less than 0.05 were considered to have significant differential expression.

#### Gene set enrichment analysis (GSEA)

GSEA was performed using significant genes ranked by log2 Fold Change resulting from DEA. The R package WebGestaltR was utilized to determine enrichment in zebrafish Gene Ontology, KEGG, Reactome, and WikiPathway databases (47). Pathways were considered enriched with an adjusted p-value < 0.05.

### Metabolic flux capacity modeling

To determine potential flux alterations of all the currently available metabolic reactions in zebrafish with CI disorders compared to healthy individuals, we applied our custom-made context-specific constraint-based metabolic modeling approach updated from previous publications (92–94). In this flux simulation, we used a recently updated zebrafish genome-scale metabolic model, Zebrafish1, containing the most extensive coverage of its metabolic network with 8,344 metabolites and 12,910 reactions. Although the zebrafish set of metabolites and reactions is smaller than other model organisms (i.e., humans), it does have sufficient representation of metabolic pathways essential to PMD such as FAO (51).

To create and optimize flux simulation based on the zebrafish model, we first maximized the reaction activity levels for certain essential pathways such as ‘Oxidative phosphorylation’, ‘Glycolysis/Gluconeogenesis’, ‘Nicotinate and nicotinamide metabolism’, ‘Fatty acid biosynthesis’, ‘Fatty acid oxidation’, ‘CoA synthesis’, and ‘CoA catabolism’. To use the Cost Optimization Reaction Dependency Assessment (CORDA), we further constrained the model to consider tissue-specific characteristics (95). To do this, we assigned confidence scores to each reaction, calculated by considering the enzyme activity level. In this computation, the activity levels were evaluated based on the gene-reaction-rule in the model using enzyme expression levels of the RNA-seq, with a total of 2,539 enzymes used in this simulation.

Finally, this custom-made flux balance analysis (FBA) iteratively optimized all metabolic reactions in the model while maximizing flux levels of their corresponding reactions regarding the aforementioned constraints under a steady state. To optimize the model, a sink reaction was incorporated to simulate ATP production, directing the flux balance analysis to prioritize pathways that maximize ATP yield within the specified constraints. Activity outside the known reactions of the metabolic model, such as the NAD/NADH ratio, are not considered. The simulation was performed using Python programming language (v. 3.6.8), Cobrapy (v. 0.14.1) (96) and Gurobi Optimizer (v. 8.1), a commercial linear programming solver (97). FBA resulting flux capacity values were compared between *ndufs2^-/-^* and *ndufs2^+/+,-/-^* larvae groups using the nonparametric Van der Waerden test in the R package matrixTests (98). Heatmaps were generated using ggplot2 (99).

### Cross-species database comparison to the *ndufs2* zebrafish model

#### Fibroblast cell lines from human LSS patients with CI gene disorders

To determine if *ndufs2^-/-^* zebrafish model has a similar transcriptomic profile to humans with a CI deficiency and is therefore a useful tool for understanding CI disorders in humans and testing therapeutic agents, gene expression of *ndufs2^-/-^*larvae (n=7) was compared to fibroblast cell lines from human LSS patients with CI subunit disorders (n=2; *NDUFS1* and *NDUFS4*). Information on patient phenotypes and genotypes can be found in **Supplemental Materials**. Fibroblast cell lines from both patients were grown in galactose media to induce metabolic stress conditions to reveal any transcriptome adaptations that are masked when cultured in glucose. The 2 CI disorder cell lines were compared to 7 human-derived fibroblast cell lines from healthy controls. RNA was extracted from human fibroblasts using the Qiagen RNeasy Plus Mini Kit (cat. # 74104) following the manufacturer’s protocol. Details on fibroblast cell culturing and RNA-Seq protocol can be found in **Supplemental Materials**.

To compare gene expression between zebrafish and fibroblast cell lines from humans affected with CI disorders, zebrafish gene symbols were converted to their human ortholog HUGO Gene Nomenclature Committee (HGNC) symbols using the R package orthogene (100). To account for the many-to-many mappings of zebrafish to human genes, with many zebrafish genes being orthologous to a single human gene, we averaged the gene expression counts of all zebrafish genes mapped to a single human gene. Pearson’s correlation analysis was performed between the log normalized average counts per million of zebrafish genes and their log normalized counts per million in humans (averaged for the two human samples). Genes with significant differential expression between *ndufs2^-/-^* and *ndufs2^+/+,+/-^* larvae were further compared to those with similar differential expression between human fibroblasts from CI disease patients (n=2) and healthy controls (n=7). Additional details on RNA-seq analysis can be found in **Supplemental Materials**.

#### gas-1(fc21) Caenorhabditis elegans

To determine if *ndufs2^-/-^* mutant zebrafish had a similar transcriptomic profile to *C. elegans* and therefore were comparable model organisms for studying CI PMD, gene expression of the *ndufs2^-/-^* larvae (n=7) was compared to previously published RNA-Seq data from *gas-1(fc21)* worms (n=4) (18). *gas-1(fc21)* mutants are a well characterized strain of mutant worms with CI deficiency due to a homozygous *Ndufs2* missense mutation (84). The processed data were further filtered using the R package WGCNA version 1.69 to remove genes with many missing entries or zero variance (90). Differential expression analysis was performed using DESeq2 (v. 1.38.3) (91) comparing all *gas-1(fc21)* mutants to healthy controls.

To compare gene expression between zebrafish and worms affected with CI disorders, all worm gene symbols were first converted to zebrafish ortholog gene symbols using the R package orthogene (100). Pearson’s correlation analysis was then performed between the log normalized average counts per million of worm genes and their log normalized counts per million in zebrafish. Genes with significant differential expression between *ndufs2^-/-^*and *ndufs2^+/+,+/-^* larvae were compared to those with similar differential expression between *gas-1(fc21)* worms and healthy N2 worms (n=4). Additional details on RNA-seq analysis can be found in **Supplemental Materials**.

### Unbiased metabolomic profiling

Twenty 7 dpf larvae were pooled and flash frozen per sample, with three samples per condition. 300 µL of ice cold 80:20 methanol:water was added to the frozen fish, then tissue was homogenized with a motorized pestle. The sample underwent three freeze-thaw cycles between liquid nitrogen and 37°C. The sample was vortexed for 1 min and centrifuged at 20,160×*g* for 15 min at 4°C. Protein quantification was performed on the supernatant using the Pierce BCA kit. A volume equivalent to 10 µg of protein was transferred to a new tube and dried in a SpeedVac. The sample was resuspended in 100 µL 80:20 acetonitrile:water. The sample was again vortexed for 1 min and centrifuged at 20,160×*g* for 15 min at 4°C. The supernatant was transferred to an LC/MS vial. Additionally, a pooled sample was made by pooling all six samples together for LC/MS data acquisition. Untargeted metabolomics were performed on a Thermo Orbitrap Exploris 480 mass spectrometer coupled to a Nexera X2 LC-30A HPLC (Shimadzu) using both positive and negative polarity modes. Metabolites are processed in a targeted manner using Sciex OS software. Differential peak analysis was conducted in R using t-tests, corrected for multiple testing using the Benjamini-Hotchberg method (101). Peaks were considered statistically significant with an adjusted p-value less than 0.05. Pathway analysis was also conducted in R using the KEGG compound pathways, as accessed through the KEGGREST R package, and the fgsea package, where pathways were considered significant with an adjusted p-value less than 0.05 (102, 103). Plots were made in R using ggplot2 (99, 104).

### Oil Red O staining for folic acid treated mutant larvae

Whole larvae Oil Red O staining at 7 dpf was performed as previously described (Lavorato). In brief, larvae were fixed in 4% paraformaldehyde (PFA) overnight at 4°C before being washed 3 times in PBS and 0.5% Tween (PBST). Next, larvae were incubated in 0.3% Oil Red O dissolved in 100% isopropyl alcohol for 15 minutes. After removing the staining solution, larvae were rinsed with PBST and incubated in 4% PFA for 10 minutes. The PFA was removed, rinsed with PBST, and larvae were stored in 50% glycerol at 4°C. Images were taken using a dissecting microscope (Olympus MVX10), an Olympus DP73 camera, and the CellSens imaging software, with qualitative analysis performed on 10 larvae per condition using ImageJ.

### Statistics

GraphPad Prism was used for statistical analysis and generating graphs for morphological and molecular characterization (version 10.0.0 for Windows, GraphPad Software, Boston, Massachusetts USA, www.graphpad.com). For phenotypic characterization, two-tailed Welch’s t-tests were performed to determine significant differences between healthy (*ndufs^+/+,+/-^*) and mutant (*ndufs2^-/-^*) zebrafish larvae (**Figure 2B & 2E – 2G**). For ETC complex activity, this test was performed pairwise for each complex (**Figure 2B**). Statistic assessment of morphology was performed using two-tailed Welch’s t-tests (**Figure 3E – 3L**). Transcriptomic analyses and visualizations were performed in the R environment (101). An unpaired, two-tailed Student’s t-test with Bonferroni correction for multiple comparisons was used for determining *ndufs2* gene expression differences (**Figure 4B**). Differential expression analysis was performed using DESeq2 (**Figure 4C**) and pathway enrichment was determined using GSEA through the R package WebGestaltR (**Figure 5**). These gene expression statistical approaches were also utilized for the cross-species transcriptomic analysis (**Figure 6B, 6C, & 6E**) and overall gene expression correlation was tested using a Pearson correlation test. Differences in the percent abundance of metabolites from untargeted metabolomics were tested in GraphPad Prism using pairwise, two-tailed Welch’s t-tests (**Figure 7B – 7E**) as well as for differences in folic acid treatment categories (**Figure 8A – 8E & 8G – 8I**). For all analyses, significance level was determined as p < 0.05 and exact p-values are provided in the Results section and figure legends.

### Study Approval

All zebrafish protocols and methods were approved by the CHOP IACUC (protocol #23-001154) and followed regulations for care and use of *D. rerio* at CHOP. Approval for study of human participants was obtained per the Children’s Hospital of Philadelphia Institutional Review Board (study #08-6177, MJF, PI).

## Data Availability

Values for all data points in graphs are reported in the Supporting Data Values file. The *D. rerio* transcriptomic data generated and analyzed for this study are available at GEO under Series ID GSE259251. Human transcriptomic data are also available under the same GEO series ID. Metabolomics data is available at the NIH Common Fund’s National Metabolomics Data Repository (NMDR) website, the Metabolomics Workbench, (https://www.metabolomicsworkbench.org) where it has been assigned Study ID ST003712 (105). *C. elegans* RNA-Seq data were previously published in Polyak et al. (2018) and are available in the Sequence Read Archive (SRA Bioproject ID PRJNA284422).

## Author Contributions

MJF conceived of, designed and oversaw performance of the study, obtained study funding, reviewed all study data and figures, and assisted in manuscript preparation. DVM performed transcriptomic data processing and analysis, statistical testing for metabolic flux modeling, and cross-species transcriptomic comparisons. DMI performed zebrafish husbandry and sample collection, lifespan testing and analysis, swim activity analysis, morphological measurements, folic acid treatment and morphological quantification. NM assisted with zebrafish swimming activity data analysis and assisted with CRISPR/Cas9 gene editing in zebrafish to create the ndufs2 deficiency zebrafish model. KK performed metabolomics analysis. CS designed and performed CRISPR/Cas9 gene editing in zebrafish to create the ndufs2 deficiency zebrafish model. SY and MK performed metabolic flux modeling. NW and DMI fixed, sectioned, stained, and imaged larval eyes. ENO performed biochemical analyses of CI function in zebrafish larvae and contributed to general methods development. DT advised and assisted with all high-throughput data and statistical analyses. DM and DI prepared figures and supplemental material for the manuscript. DVM, DMI, VEA, and MJF drafted the manuscript. All authors reviewed and approved the final manuscript.

## Acknowledgements

We are grateful to the individuals with mitochondrial complex I disease and families who consented to allow their cells to be studied in this research program. This work was funded in part by the National Institutes of Health (NIH, R35-GM134863, M.J.F., PI; T32-NS007413; T32HG009495-06). Metabolomics Workbench is supported by NIH grant U2C-DK119886 and OT2-OD030544 grants. The content is solely the responsibility of the authors and does not necessarily represent the official views of the NIH.

## Notes

https://zebrafish.org/

https://zfin.org

https://www.metabolomicsworkbench.org/data/DRCCMetadata.php?Mode=Study&StudyID=ST003712

### Competing Interest Statement

MJF is inventor of US patent 12,011,452 B2 issued Jun 18, 2024, Compositions and Methods for Treatment of Mitochondrial Respiratory Chain Dysfunction and Other Mitochondrial Disorders. MJF is engaged with companies involved in mitochondrial disease therapeutic preclinical and/or clinical-stage development. MJF is co-founder of Rarefy Therapeutics; an advisory board member with equity interest in RiboNova Inc.; a scientific advisory board member and paid consultant with Khondrion and Larimar Therapeutics; has been a paid consultant for Astellas (formerly Mitobridge), Ajinomoto Cambrooke, Casma Therapeutics, Cyclerion Therapeutics, Epirium Bio (formerly Cardero Therapeutics), HealthCap VII Advisor AB, Imel Therapeutics, Mayflower, Inc., Primera Therapeutics, Inc., Minovia 2 Therapeutics, Mission Therapeutics, NeuroVive Pharmaceutical AB, Precision Biosciences, Reneo Therapeutics, Saol Therapeutics, Stealth BioTherapeutics, incere Bio, and Zogenix; and/or has been a sponsored research collaborator for Aadi Bioscience, Adjuvia Therapeutics, Astellas, Cyclerion Therapeutics, Epirium Bio, Imel Therapeutics, Khondrion, Merck, Minovia Therapeutics, Mission Therapeutics, NeuroVive Pharmaceutical AB, Precision Biosciences, PTC Therapeutics, Raptor Pharmaceuticals, REATA Inc., Reneo Therapeutics, RiboNova, Saol Therapeutics, Standigm, Stealth BioTherapeutics, and Thiogenesis. MJF also has received royalties from Elsevier and speaker fees from Agios Pharmaceuticals, GenoMind, and educational honorarium from PlatformQ. None of the other authors have relevant conflicts of interest to declare.

