## Supplemental Material for "*ndufs2^-/-^* zebrafish have impaired survival, neuromuscular activity, morphology, and one-carbon metabolism treatable with folic acid"

### SUPPLEMENTAL MATERIALS.

#### SUPPLEMENTAL METHODS.

##### $NAD^+$ and NADH Measurement by HPLC Analysis in zebrafish larvae.

Zebrafish larvae at 7 dpf were collected and washed two times with E3 buffer, and flash-frozen in aliquots of 20 larvae per tube, with 3-4 biological replicates per condition. Samples were prepared by established methods using perchloric acid (PCA) extraction for $NAD^+$ and ammonium acetate/acetonitrile/NaOH extraction for NADH (PMID: 25411251). Separation of $NAD^+$ and NADH was performed with modifications using an YMC-Pack ODS-A column (5 µm, 4.6 × 250 mm) preceded by a guard column at 50 °C. Flow rate was set at 0.4 ml/min. The mobile phase was initially 100% mobile phase A (0.1 m potassium phosphate buffer, pH 6.0). Methanol was linearly increased with mobile phase B (0.1 m potassium phosphate buffer, pH 6.0 containing 30% methanol) to 50% over 15 min. The column was washed after each separation by increasing mobile phase B to 100% for 5 min. UV absorbance was monitored at 260 and 340 nm with a Shimadzu SPD-M20A. The column was washed after each separation by increasing mobile phase B to 100% for 3 min. Pertinent peak areas were integrated by the LabSolution software from Shimadzu, quantified using standard curves, and normalized to protein concentration. GraphPad Prism was used for generating graphs and statistical analysis was done by two-tailed, unpaired Student's t-test. Three or four biological replicate experiments were performed for each condition.

##### Transcriptomics

*Sample preparation*

Qiagen RNeasy Plus Mini Kit (cat. # 74134) was used for the RNA extraction from zebrafish larvae samples, following the manufacturer's protocol. Briefly, the fish were homogenized using a tissue grinder with 350 µL RLT buffer and 20 ng carrier RNA. After centrifugation, the lysate was passed through a gDNA eliminator column and the flow through was mixed with 70% ethanol. The sample was then passed through an RNeasy MinElute column and washed with RW1 buffer, RPE buffer, and 80% ethanol. The RNA was then eluted in 14 µL of RNase-free water. The concentration was determined with a Nanodrop® (Thermo Scientific) and a Tapestation Bioanalyzer (Agilent) was used to assess RNA quality. Libraries were generated using a

Sciclone® NGSx Workstation (PerkinElmer). PolyA-selected RNA sequencing was performed using an Illumina Novaseq6000 to obtain 100 bp paired-end reads.

*Alignment & Quantification*

RNA-Seq data were preprocessed using the nf-core RNAseq pipeline version 3.9 (72). FastQC version 0.11.9 was used to assess quality of the reads (73). Trim Galore! version 0.6.7 was then used for adapter and quality trimming on the raw reads (74). Reads were aligned to GRCz11 (danRer11) using STAR version 2.7.10a (75) and quantified using RSEM version 1.3.1 (76). Quantification was based on the Lawson Lab Transcriptome Annotation v4.3.2 (47). Duplicates were removed using picard MarkDuplicates version 2.27.4-snapshot (77).

**Cross-species database comparison to the *ndufs2* zebrafish model**

*Fibroblast cell lines from human LSS patients with CI gene disorders*

Patient descriptions: The first patient exhibited frequent vomiting, leukodystrophy, developmental delay, feeding and swallowing difficulties, GERD, increased oropharyngeal secretions, thrombocytopenia, macrocephaly, diarrhea, hypotonia, and fatigue. Genetic testing confirmed this patient was a compound heterozygote with a deletion and missense mutation in the NDUFS1 gene (c.592_594delACA:p.T198del and c.1727G>A:p.G576E). The second patient's phenotype included chronic respiratory failure, hypertension, leukodystrophy, renal tubular acidosis, hypercortisolism, and bilateral hearing loss. This patient's LSS was confirmed to result from a homozygous premature stop in the NDUSF4 gene (c.377_384delTAACCTTC:p.L126QfsX3).

Human fibroblast cell culturing: Human fibroblast cell lines derived from skin biopsies were grown in DMEM supplemented with a low level of glucose and reached 80% confluence before being washed and replated with 10 mM galactose. Cells grown in galactose for 24 hours were then collected for sequencing, as galactose stresses cells by requiring OXPHOS function and unmasking transcriptional alterations to mitochondrial dysfunction that may be 'treated' with glucose.

RNA-Seq methodology: RNA was extracted from human fibroblasts using the Qiagen RNeasy Plus Mini Kit (cat. # 74104) following the manufacturer's protocol. Briefly, 200 µL of chloroform was added to 1 mL TRIzollysed cells (Thermo Fisher, cat# 15596026). After centrifugation, the lysate was mixed with 70% ethanol and passed through an RNAeasy MinElute column then washed with 700 µL of RW1 buffer and twice with 500 µL RPE buffer. The RNA was then eluted with 50 µL of RNase-free water. The concentration was determined with a Nanodrop® (Thermo Scientific) and a Tapestation Bioanalyzer (Agilent) was used to assess RNA quality. Libraries were generated using the RNA-seq NETFLEX Rapid Directional kit version 1.0 from PerkinElmer. PolyA-selected RNA sequencing was performed using an Illumina Novaseq6000 to obtain 100 bp paired-end reads. RNA-seq data were then aligned using STAR (75) against GRCH38 (hg38) version of the human genome. Raw read counts were quantified using RSEM (76). Gene expression data were filtered using the R package WGCNA version 1.69 to remove genes with many missing entries or zero variance (78). For cross-species comparisons of gene-to-gene expression, gene expression was averaged between the two CI patients and normalized using counts per million (cpm). Differential expression analysis was performed using DESeq2 (v. 1.38.3) (79) comparing all CI patients (n=2) to healthy controls (n=7).

**Supplemental Figures & Tables.**

```
C. elegans  MLGRKIAGTCLRANV----PSVAATSSTPATQTRNSHTIWYPDAKFERQFKTGGTLGKLW    56
D. rerio    ------------------------------------------------------------    0
H. sapiens  --MAALRALCGFRGVAAQVLRPGAG-VRLPIQPSRGVRQWQPDVEWAQQFGGAVMY----    53

C. elegans  MSERVSDFDEKIGLDKLEKLAYSDPVMSDNYSGKQREKNLENMILNFGPQHPAAHGVLRL    116
D. rerio    --------------------------------------------------------MIRI    4
H. sapiens  PS--------------KETAHWKPPPWND--VDPPKDTIVKNITLNFGPQHPAAHGVLRL    97
                                                                    ::*:

C. elegans  VLKLEGEVIIKAIPHIGLLHRATEKLIEHKTYTQALPYFDRLDYVSMMCNEQAWSLAVEK    176
D. rerio    HQHRRICPTSPSTSGLSTLHRGTEKLIEYKTYLQALPYFDRLDYVSTMCNEQAYSLAVEK    64
H. sapiens  VMELSGEMVRKCDPHIGLLHRGTEKLIEYKTYLQALPYFDRLDYVSMMCNEQAYSLAVEK    157
              .        .   :. ***.******:*** ************* ******:******

C. elegans  LLGIDIPTRAKYIRTLMGELTRIQNHIMGITTHALDVGAMTPFFWMFEEREKLFEFSERV    236
D. rerio    LLNIQAPPRAQWIRVLFGEMTRIMNHIMGITTHALDIGAMTPFFWMFEEREKLFEFYERV    124
H. sapiens  LLNIRPPPRAQWIRVLFGEITRLLNHIMAVTTHALDLGAMTPFFWLFEEREKMFEFYERV    217
            **.*  * **::**.*:**:**: ****.:******:********:******:*** ***

C. elegans  SGARMHANYVRPGGVAWDLPIGLMDDIYDWAIKFPERIDELEDMLTENRIWKARTIDIGL    296
D. rerio    SGARMHAAYVRPGGVHQDMPLGLMDDIYEWCKNFSIRIDEVEEMLTNNRIWKNRTVGIGV    184
H. sapiens  SGARMHAAYIRPGGVHQDLPLGLMDDIYQFSKNFSLRLDELEELLTNNRIWRNRTIDIGV    277
            ******* *:*****  *:*:*******::. :*  *:**:*::**:****: **:.**:

C. elegans  VSAADALNWGFSGVMVRGSGIKQDVRKTEPYDAYADMEFDVPIGTKGDCYDRYLCRIEEM    356
D. rerio    IGAEEALNYGFSGVMLRGSGIKWDLRKSQPYDKYDEVEFDMAIGTNGDCYDRYLCRVEEM    244
H. sapiens  VTAEEALNYGFSGVMLRGSGIQWDLRKTQPYDVYDQVEFDVPVGSRGDCYDRYLCRVEEM    337
            : * :***:******:*****: *:**::*** * ::***: :*:.**********:***

C. elegans  RQSLNIVHQCLNKMPAGEIKVDDHKVVPPKRAEMKENMESLIHHFKFFTEGFQVPPGATY    416
D. rerio    RQSLRIMHQCLNKMPPGEIKVDDAKIAPPKRSEMKTSMESLIHHFKLYTEGYQVPPGATY    304
H. sapiens  RQSLRIIAQCLNKMPPGEIKVDDAKVSPPKRAEMKTSMESLIHHFKLYTEGYQVPPGATY    397
            ****.*: ******* ******* *: ****:*** .*********::***:********

C. elegans  VPIEAPKGEFGVYLVADGTGKPYRCFIRAPGFAHLAAIHDVCYMSLIADIVAVIGTMDIV    476
D. rerio    TAIEAPKGEFGVYLVSDGSSRPYRCKIKAPGFAHLAGLDRMSQGHMLADVVAIIGTQDIV    364
H. sapiens  TAIEAPKGEFGVYLVSDGSSRPYRCKIKAPGFAHLAGLDKMSKGHMLADVVAIIGTQDIV    457
            . *************:**:.:**** *:********.:. :.   ::**:**:*** ***

C. elegans  FGEVDR    482
D. rerio    FGEVDR    370
H. sapiens  FGEVDR    463
            ******
```

**Supplemental Figure S1**: NDUFS2 is highly conserved in humans, *D. rerio*, and *C. elegans*. Protein alignment is shown where below the symbol of a star indicates identity, a colon indicates a highly similar amino acid, a period indicates a semi-conserved amino acid, and no symbol is no conservation across all three species. The amino acid identity between zebrafish and humans is 87%.

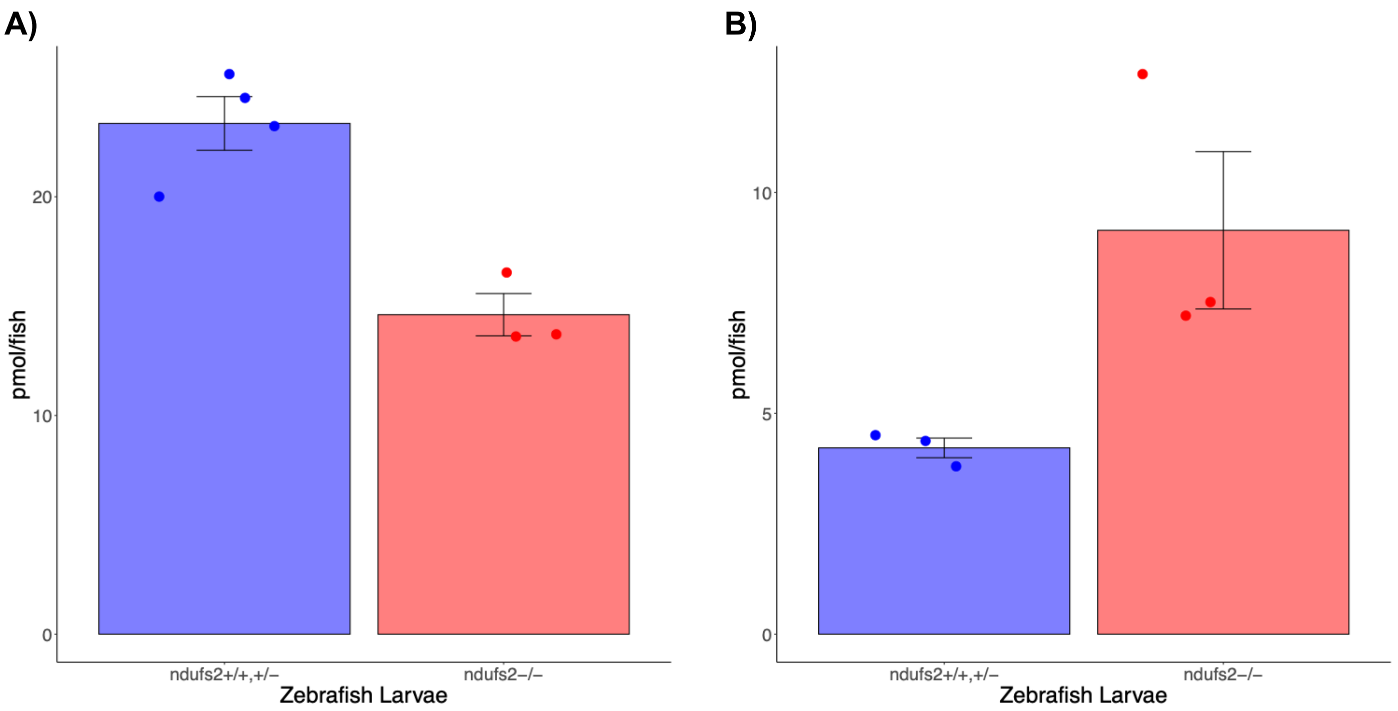


**Supplemental Figure S2. NAD$^+$ and NADH Measurement by HPLC. A)** NAD$^+$ measurements in samples of healthy *ndufs2*$^{+/+,+/-}$ larvae (n=4) compared with mutant *ndufs2*$^{-/-}$ larvae (n=3). Each sample contains 20 zebrafish larvae. Mutant zebrafish larvae had significantly lower pmol/fish (unpaired, two-sided Student's t-test, $p = 0.002$). **B)** NADH measurements in samples of healthy *ndufs2*$^{+/+,+/-}$ larvae (n=3) compared with mutant *ndufs2*$^{-/-}$ larvae (n=3). Each sample contains 20 zebrafish larvae. Mutant zebrafish larvae had marginally significantly higher pmol/fish (unpaired, two-sided Student's t-test, $p = 0.107$).

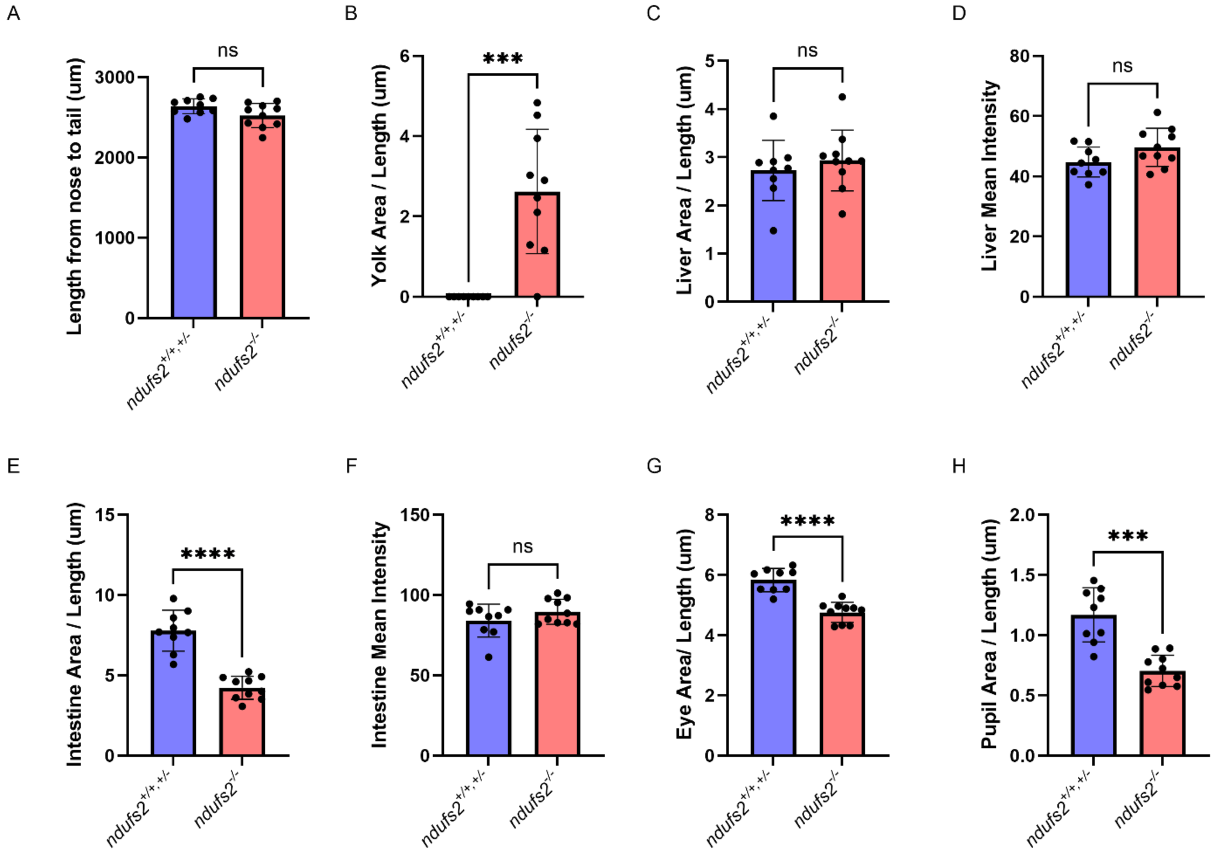


**Supplemental Figure S3**: **7 dpf *ndufs2$^{-/-}$* larvae morphological parameter measurements.** Organs were quantified at 7 dpf: A, length from nose to tail; B, yolk area; C, liver area; D, liver mean intensity; E, intestine area; F, intestine mean intensity; G, eye area; H, pupil area. Differences between larval groups were identified using a two-tailed Student’s t-test: ***, $P < 0.001$; ****, $P < 0.0001$; ns, no statistically significant difference. Each point represents a measurement from one larva. The bars represent the average and the error bars represent standard deviation.

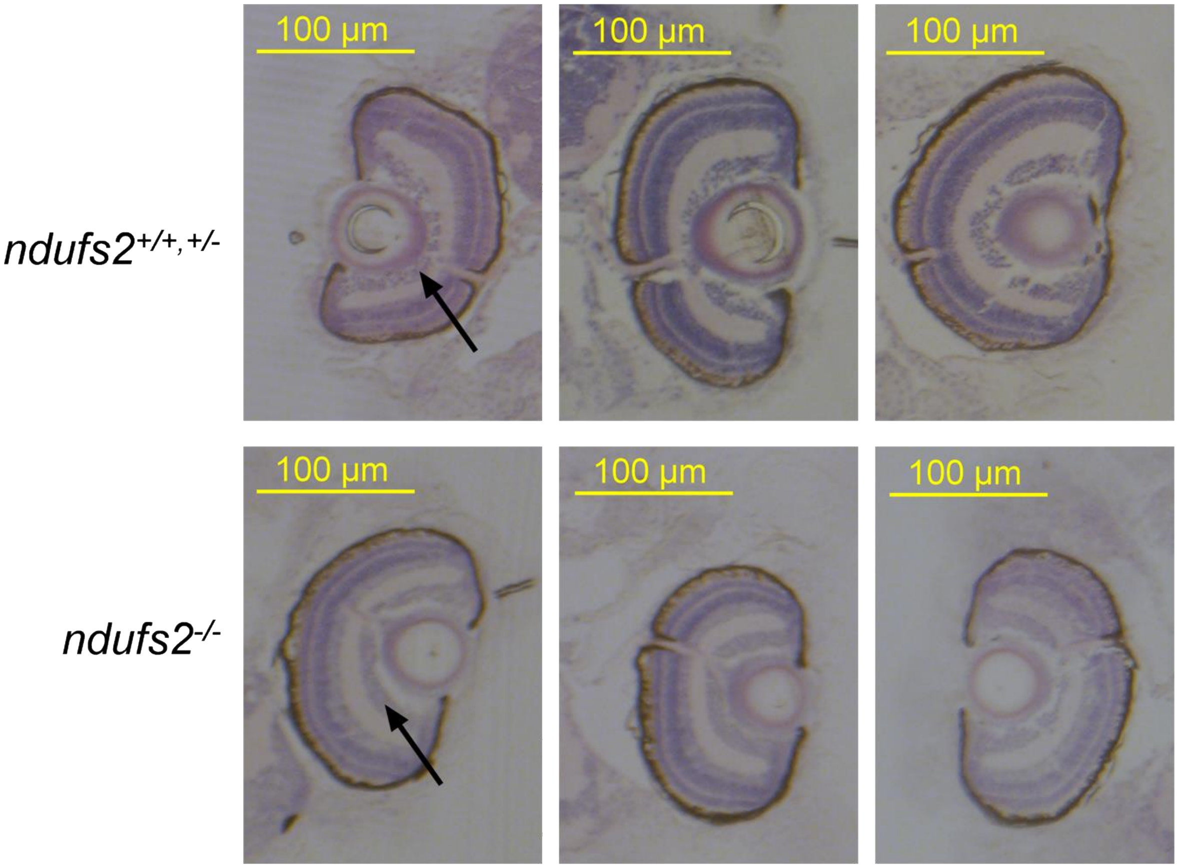


**Supplemental Figure S4: Retinal ganglion cell layer differs in *ndufs2*$^{-/-}$ larvae.** Hematoxylin and eosin stain of 5 µm sections of 7 dpf larvae. Black arrow indicates the ganglion cell layer.

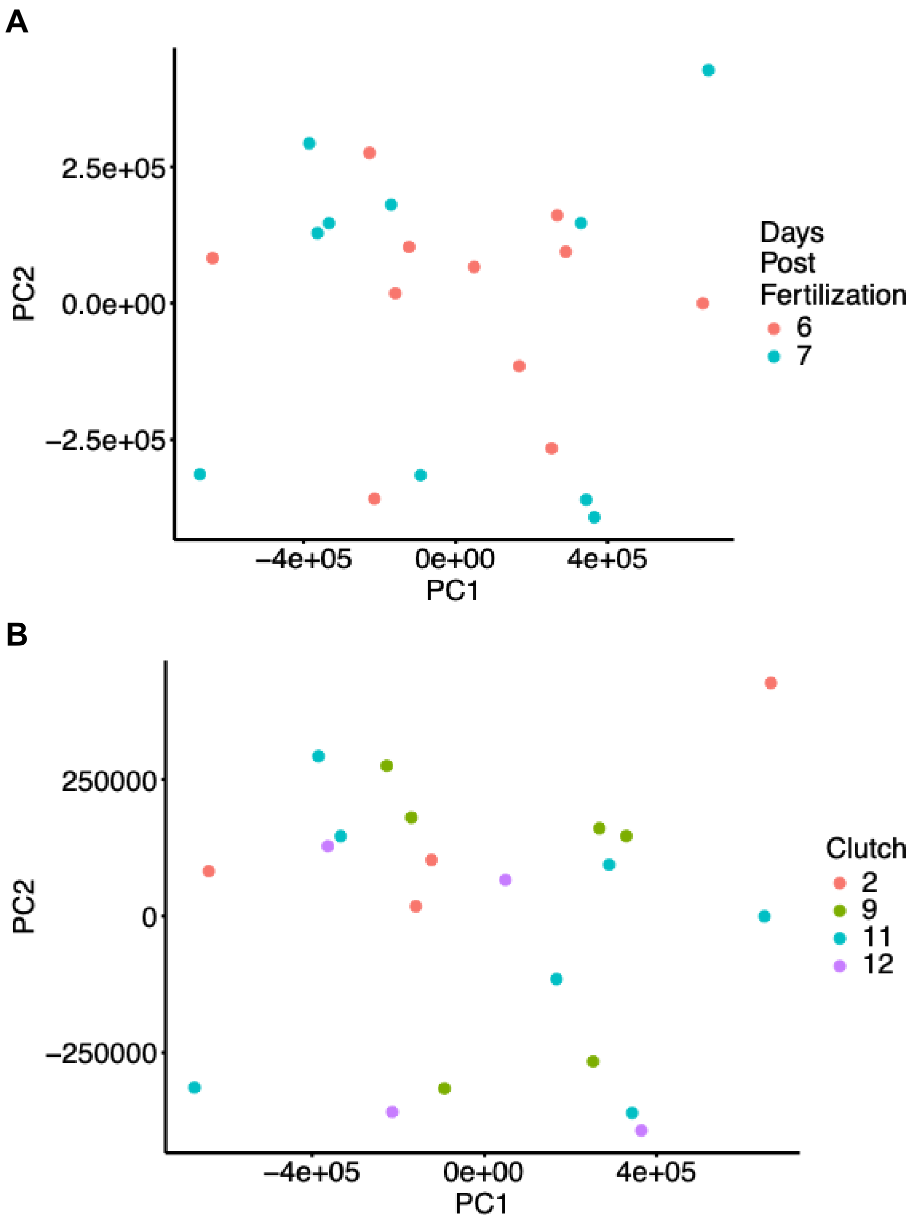


**Supplemental Figure S5: Principal component analysis (PCA) of zebrafish transcriptomic data with batches represented.** A) Zebrafish larvae from all lines (AB, *ndufs2*$^{+/+,+/-}$, and *ndufs2*$^{-/-}$) are marked by days post fertilization (dpf) at which the sample was harvested, either 6 or 7 dpf. B) Zebrafish larvae from all lines (AB, *ndufs2*$^{+/+,+/-}$, and *ndufs2*$^{-/-}$) are marked by the fish clutch from which the sample was harvested.

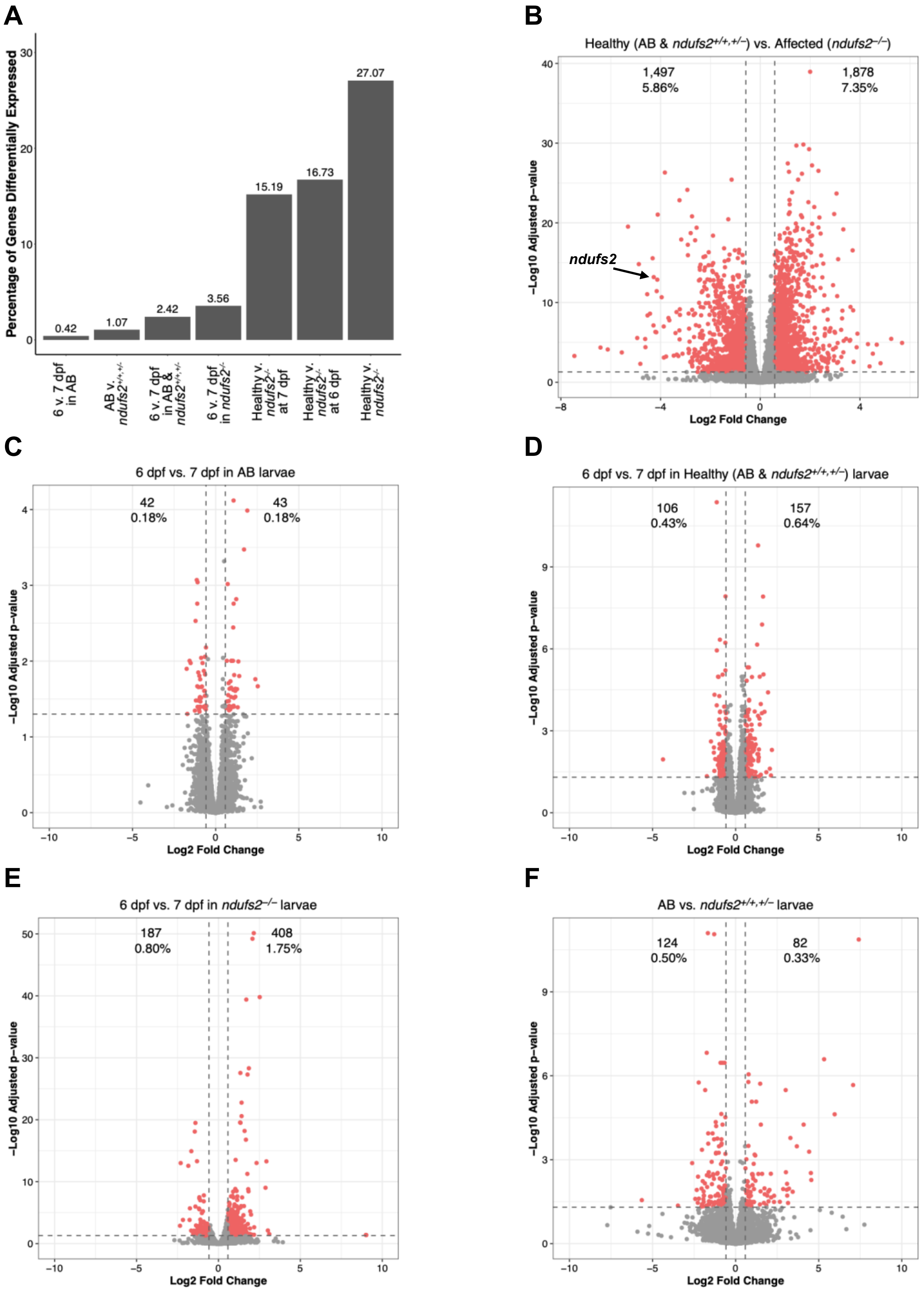
A
Percentage of Genes Differentially Expressed
0.42
1.07
2.42
3.56
15.19
16.73
27.07
6 v. 7 dpf in AB
AB v. ndufs2+/+,+/-
6 v. 7 dpf in AB & ndufs2+/+,+/-
6 v. 7 dpf in ndufs2-/-
Healthy v. ndufs2-/- at 7 dpf
Healthy v. ndufs2-/- at 6 dpf
Healthy v. ndufs2-/-
B
Healthy (AB & ndufs2+/+,+/–) vs. Affected (ndufs2–/–)
1,497
5.86%
1,878
7.35%
ndufs2
–Log10 Adjusted p–value
Log2 Fold Change
C
6 dpf vs. 7 dpf in AB larvae
42
0.18%
43
0.18%
D
6 dpf vs. 7 dpf in Healthy (AB & ndufs2+/+,+/–) larvae
106
0.43%
157
0.64%
E
6 dpf vs. 7 dpf in ndufs2–/– larvae
187
0.80%
408
1.75%
F
AB vs. ndufs2+/+,+/– larvae
124
0.50%
82
0.33%

**G**

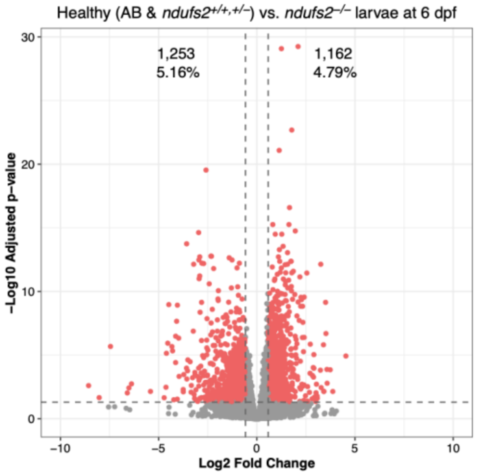


**H**

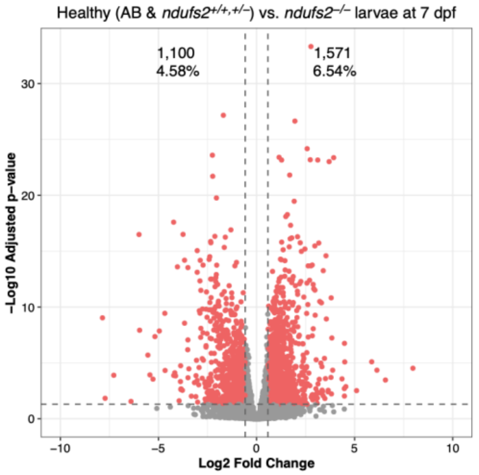


**Supplemental Figure S6**: **Differential expression analysis of RNA-Seq data between zebrafish larval batches.** A) Summary of the number of genes with significant differential expression (calculated with DESEq2) for each comparison between batches of genotype and dpf (across x-axis). Batches include from left to right: AB wild-type zebrafish at 6 dpf compared to AB wild-type zebrafish at 7 dpf; AB wild-type zebrafish at both 6 and 7 dpf compared to in cross zebrafish siblings (genotype *ndufs2*$^{+/+,+/-}$) at both 6 and 7 dpf; All healthy zebrafish including AB wild-type and healthy in cross siblings (genotype *ndufs2*$^{+/+,+/-}$) at 6 dpf compared to the same group at 7 dpf; Mutant *ndufs2*$^{-/-}$ zebrafish at 6 dpf compared to the same group at 7 dpf; All healthy zebrafish from 6 dpf including AB wild-type and healthy in cross siblings (genotype *ndufs2*$^{+/+,+/-}$) compared to mutant *ndufs2*$^{-/-}$ zebrafish at 6 dpf;  All healthy zebrafish from 7 dpf including AB wild-type and healthy in cross siblings (genotype *ndufs2*$^{+/+,+/-}$) compared to mutant *ndufs2*$^{-/-}$ zebrafish at 7 dpf;  All healthy zebrafish from 6 and 7 dpf including AB wild-type and healthy in cross siblings (genotype *ndufs2*$^{+/+,+/-}$) compared to all mutant *ndufs2*$^{-/-}$ zebrafish at 6 and 7 dpf. Number of differentially expressed genes is given as a proportion of the total number of genes tested for each comparison and indicated at the top of each bar. B) Volcano plot of differential gene expression between *ndufs2*$^{-/-}$ and *ndufs2*$^{+/+,+/-}$ larvae. C) Volcano plot of differential gene expression between 6 dpf and 7 dpf in AB zebrafish larvae. D) Volcano plot of differential gene expression between 6 dpf and 7 dpf in all healthy zebrafish larvae (AB and *ndufs2*$^{+/-}$ larvae). E) Volcano plot of differential gene expression between 6 dpf and 7 dpf in *ndufs2*$^{-/-}$ zebrafish larvae. F) Volcano plot of differential gene expression between control zebrafish lines: AB vs. *ndufs2*$^{+/+,+/-}$ larvae. G) Volcano plot of differential gene expression between all healthy (AB and *ndufs2*$^{+/+,+/-}$) and *ndufs2*$^{-/-}$ zebrafish larvae at 6 dpf only. H) Volcano plot of differential gene expression between all healthy (AB and *ndufs2*$^{+/+,+/-}$) and *ndufs2*$^{-/-}$ zebrafish larvae at 7 dpf only. For all volcano plots, genes with significantly different expression (adjusted p-value < 0.05) and large log2 Fold Change (log2FC > 0.58) are highlighted in red.

**Supplemental Figure S7: Metabolic flux balance analysis of *ndufs2*$^{-/-}$ larvae transcriptomic data**. Biochemical flux alterations of *ndufs2*$^{-/-}$ larvae were modeled in the pentose phosphate pathway, glycolysis, and TCA cycle. The subsystem is divided into components located in the cytosolic and mitochondrion compartments, as designated by the black dotted lines. Biochemical reaction arrows represent the log2 Fold Change between predicted flux values of *ndufs2*$^{-/-}$ larvae relative to *ndufs2*$^{+/+,+/-}$ larvae. Arrow highlights indicate directionality of the biochemical reaction in *ndufs2*$^{-/-}$ compared to *ndufs2*$^{+/+,+/-}$ larvae: red, blue, and gray highlights, respectively, indicate reactions with increased, decreased, or no alteration in flux in. No data were available to predict flux for reactions conveyed by black arrows. Metabolites are indicated in lowercase black text. Reaction IDs from the BiGG Knowledgebase are displayed in capital blue letters only for reactions with differences between groups. Metabolic flux values and statistical analysis results indicating inter-group differences are detailed in **Supplementary Table S3**.

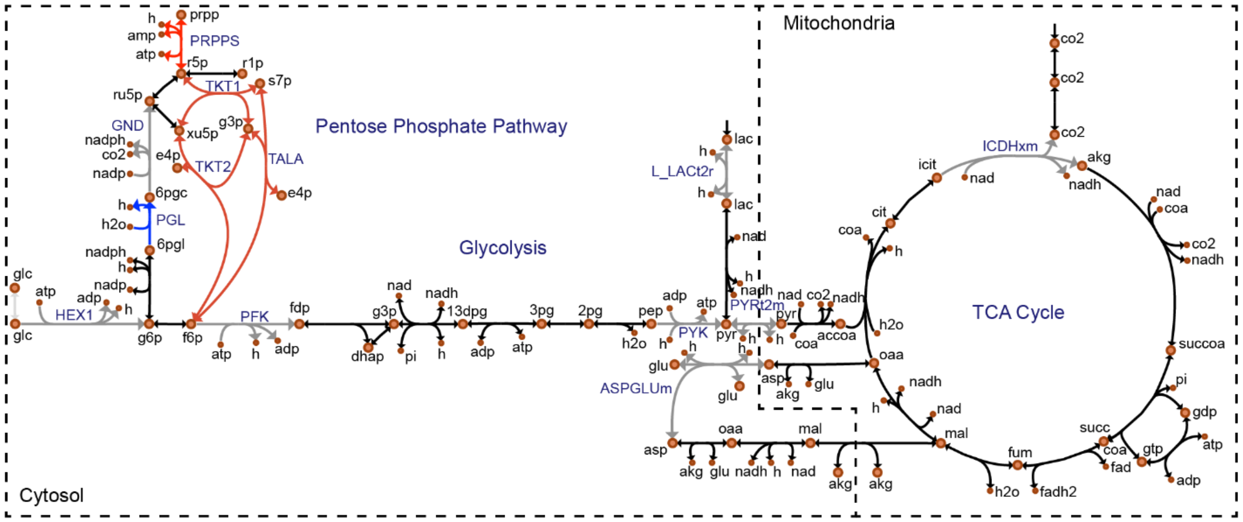

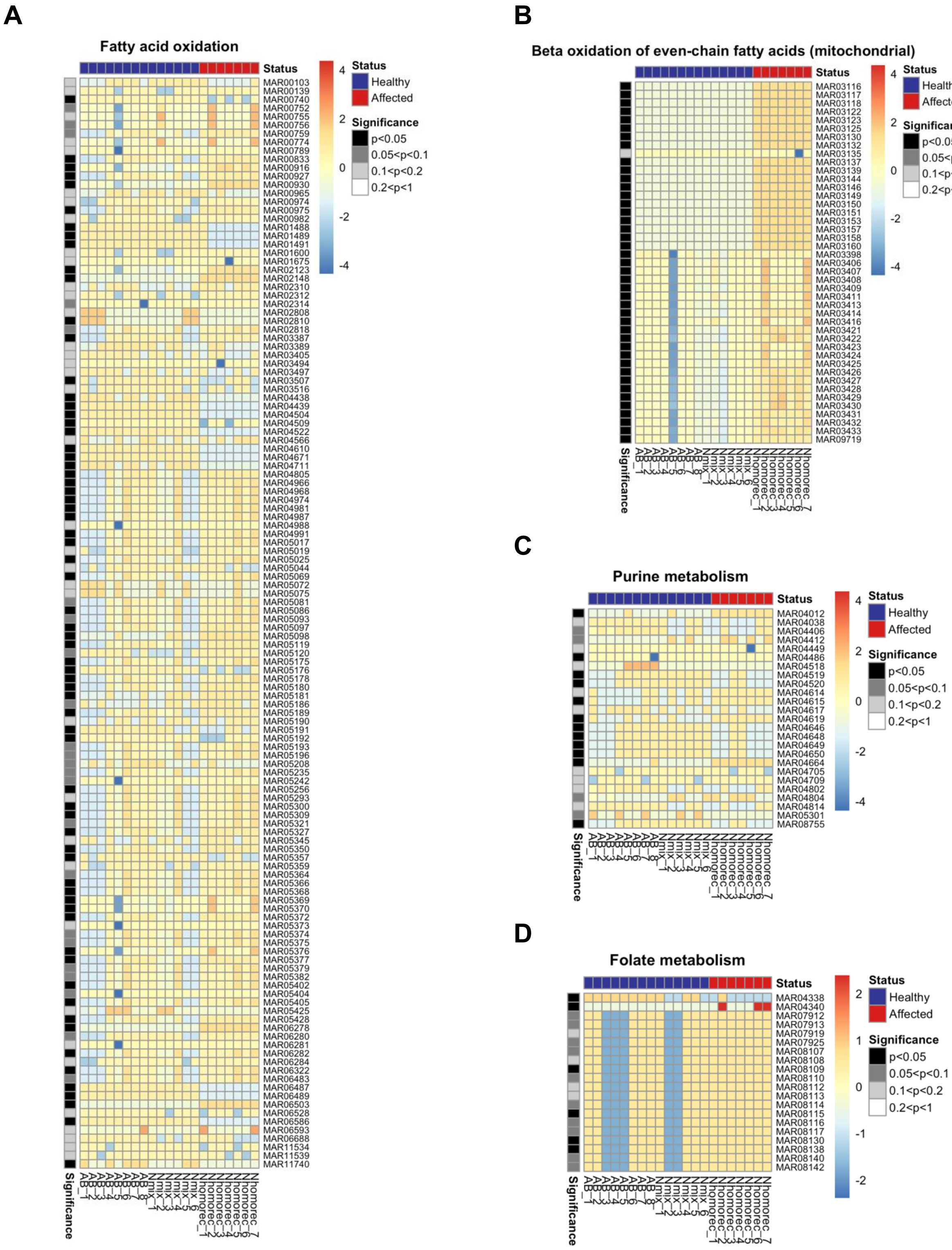


**Supplemental Figure S8**: Heatmaps showing flux values of metabolic atlas reaction IDs (MAR#s, https://metabolicatlas.org/explore/Human-GEM/gem-browser) from the BiGG knowledgebase (rows) for each zebrafish sample (columns) categorized as affected (*ndufs2*$^{-/-}$ larvae) in blue or healthy (*ndufs2*$^{+/+,+/-}$) in red from reactions in the A) Fatty acid oxidation, B) Beta oxidation of even-chained fatty acids (mitochondrial) subsystems, C) Purine biosynthesis, and D) Folate metabolism. P-values were calculated using the Van der Waerden test and are shown for each row as indicated by the grey scale.

A

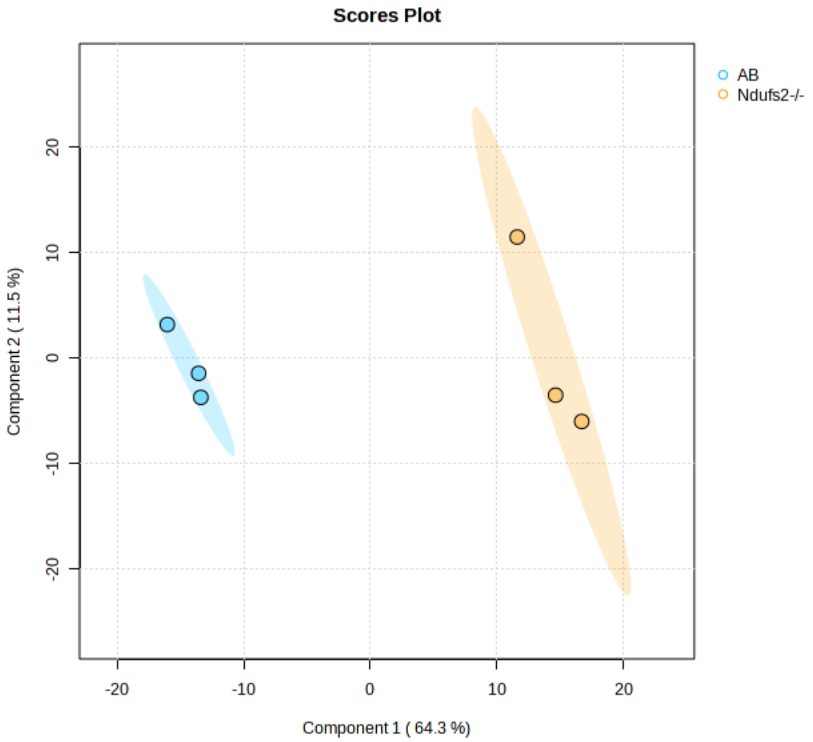


B

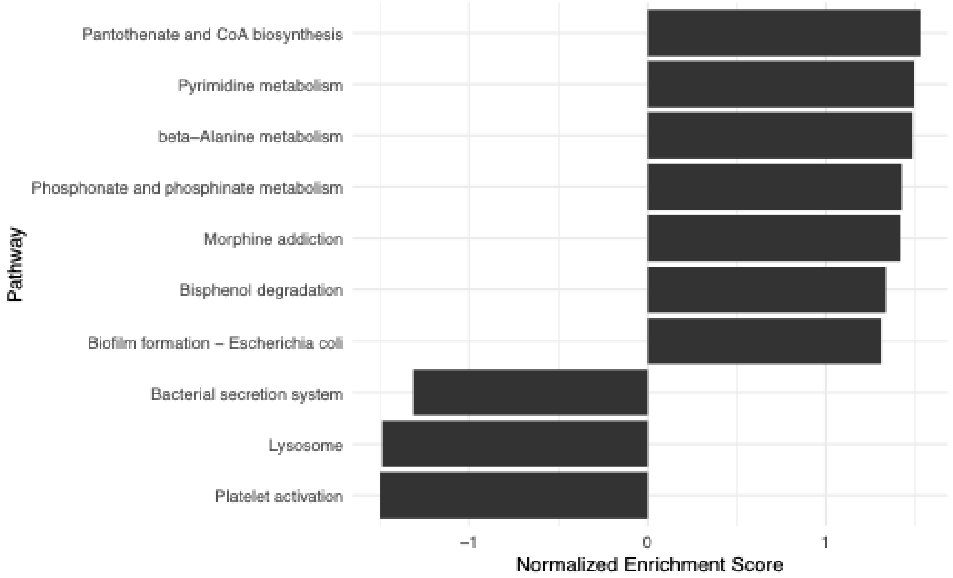


**Supplemental Figure S9. A)** Partial least squares-discriminant analysis (PLS-DA) from the metabolomics dataset for AB (n=3) and *ndufs2*$^{-/-}$ larvae (n=3). **B)** Pathway enrichment analysis of metabolites in *ndufs2*$^{-/-}$ larvae compared to AB larvae.

| Sample ID | Zebrafish Line | Days Post Fertilization (DPF) | Clutch | Status |
|---|---|---|---|---|
| 1 | AB | 6 | A | Wild-type |
| 2 | AB | 6 | A | Wild-type |
| 3 | AB | 6 | A | Wild-type |
| 4 | AB | 6 | B | Wild-type |
| 5 | AB | 6 | C | Wild-type |
| 6 | AB | 7 | B | Wild-type |
| 7 | AB | 7 | C | Wild-type |
| 8 | AB | 7 | A | Wild-type |
| 9 | *ndufs2*$^{+/+,+/-}$ | 6 | D | Wild-type |
| 10 | *ndufs2*$^{+/+,+/-}$ | 6 | E | Wild-type |
| 11 | *ndufs2*$^{+/+,+/-}$ | 6 | F | Wild-type |
| 12 | *ndufs2*$^{+/+,+/-}$ | 7 | D | Wild-type |
| 13 | *ndufs2*$^{+/+,+/-}$ | 7 | E | Wild-type |
| 14 | *ndufs2*$^{+/+,+/-}$ | 7 | F | Wild-type |
| 15 | *ndufs2*$^{-/-}$ | 6 | D | Affected |
| 16 | *ndufs2*$^{-/-}$ | 6 | E | Affected |
| 17 | *ndufs2*$^{-/-}$ | 6 | F | Affected |
| 18 | *ndufs2*$^{-/-}$ | 7 | D | Affected |
| 19 | *ndufs2*$^{-/-}$ | 7 | E | Affected |
| 20 | *ndufs2*$^{-/-}$ | 7 | F | Affected |
| 21 | *ndufs2*$^{-/-}$ | 7 | E | Affected |

**Supplemental Table S1: Detailed sample information for all zebrafish larvae studied by RNA-Seq analysis, including genotype, dpf, and clutch.** "*ndufs2*$^{+/+,+/-}$" indicates wild-type mixed genotype siblings of *ndufs2*$^{-/-}$ mutants from *ndufs2*$^{+/-}$ heterozygous in-cross. "AB" conveys unrelated, age-matching wild-type control.

| Reaction ID | Reaction Name | Average *ndufs2*$^{-/-}$ Flux | Average *ndufs2*$^{+/+,+/-}$ Flux | Log2 Fold Change |
|---|---|---|---|---|
| **Pentose Phosphate Pathway** | | | | |
| PRPPS | Phosphoribosylpyrophosphate synthetase | 1000 | 929 | -0.107 |
| TKT1 | Transketolase | 999 | 952 | -0.0704 |
| GND | Phosphogluconate dehydrogenase | 1000 | 1000 | 0 |
| TALA | Transaldolase | 1000 | 952 | -0.0705 |
| TKT2 | Transketolase | 1000 | 952 | -0.0711 |
| PGL | 6-Phosphogluconolactonase | 286 | 500 | 0.807 |
| HEX1 | Hexokinase (D-glucose:ATP) | 1000 | 1000 | 0 |
| PFK | Phosphofructokinase | 1000 | 1000 | 0 |
| PYK | Pyruvate kinase | 1000 | 1000 | 0 |
| ASPGLUm | ASPGLUm | 1000 | 1000 | 0 |
| L_LACt2r | Lactate reversible transport via proton symport | 1000 | 1000 | 0 |
| PYRt2m | Pyruvate mitochondrial transport via proton symport | 1000 | 1000 | 0 |
| ICDHxm | Isocitrate dehydrogenase NAD | 1000 | 1000 | 0 |
| **Glycolysis** | | | | |
| L_LACt2r | L lactate reversible transport via proton symport | 1000 | 1000 | 0 |
| PYK | Pyruvate kinase | 1000 | 1000 | 0 |
| PYRt2m | Pyruvate mitochondrial transport via proton symport | 1000 | 1000 | 0 |
| ASPGLUm | Aspartate-glutamate mitochondrial shuttle | 1000 | 1000 | 0 |
| **TCA Cycle** | | | | |
| ICDHxm | Isocitrate dehydrogenase NAD | 1000 | 1000 | 0 |

**Supplemental Table S2.** Metabolic flux calculation of reactions in the pathways shown in **Figure 6**, including the pentose phosphate pathways, TCA cycle, and Glycolysis. Average flux values were calculated using flux balance analysis of transcriptomic data. Reaction IDs from the BiGG Knowledgebase and their full names are shown along with predicted metabolic flux values averaged across *ndufs2*$^{-/-}$ larvae and *ndufs2*$^{+/+,+/-}$ larvae as well as log2 Fold Change between the two groups.
